# Harnessing Molecular Motors for Nanoscale Pulldown in Live Cells

**DOI:** 10.1101/053744

**Authors:** Jonathan E. Bird, Melanie Barzik, Meghan C. Drummond, Daniel C. Sutton, Spencer M. Goodman, Eva L. Morozko, Stacey M. Cole, Jennifer Skidmore, Diana Syam, Elizabeth A. Wilson, Tracy Fitzgerald, Atteeq U. Rehman, Donna M. Martin, Erich T. Boger, Inna A. Belyantseva, Thomas B. Friedman

**Author notes:** Equal Contribution (Jonathan E. Bird, Melanie Barzik). To whom correspondence should be addressed: Jonathan E. Bird, Ph.D., National Institute on Deafness and Other Communication Disorders, Building 35, Room 1F-145, 9000 Rockville Pike, Bethesda, MD 20892, USA., E-mail: Correspondence.

## Abstract

Protein-protein interactions (PPIs) regulate signal transduction and cellular behavior, yet studying PPIs within live cells remains fundamentally challenging. We have miniaturized the affinity pulldown, a gold-standard PPI interrogation technique, for use within live cells. Our assay hijacks endogenous myosin motors to forcibly traffic, or pulldown, macromolecular complexes within the native cytosolic environment. Macromolecules captured by nanoscale pulldown (NanoSPD) are optically interrogated *in situ* by tagging individual protein components. Critically, continuous motor trafficking concentrates query complexes into nanoscopic subcellular compartments, providing fluorescence enhancement and allowing nanoscale pulldowns to be visualized and quantified by standard microscopy. Nanoscale pulldown is compatible with nuclear, membrane-associated and cytoplasmic proteins and can investigate functional effects of protein truncations or amino acid substitutions. Moreover, binding hierarchies in larger complexes can be quickly examined within the natural cytosol, making nanoscale pulldown a powerful new optical platform for quantitative high-content screening of known and novel PPIs that act within macromolecular assemblies.

## INTRODUCTION

The identification of protein-protein interactions (PPI) within macromolecular complexes is a powerful approach to understanding cellular biology in normal and disease states. Many methodologies exist to discover the composition and underlying PPIs that govern assembly and turnover of these complexes. These include yeast and mammalian two-hybrid assays and affinity purification coupled with mass spectrometry (Fields et al., 1989, Gingras et al., 2007, Lemmens et al., 2015, Luo et al., 1997, Shioda et al., 2000). Because these methodologies vary in their degree of sensitivity and specificity, no single approach has been shown to eliminate false-positive and false-negatives to detect the full landscape of PPIs (Braun et al., 2009, Chen et al., 2010). Replication through independent orthogonal assays is thus critical to build confidence in the presence or absence of a putative PPI.

Co-immunoprecipitation (IP) and affinity pulldown (AP) assays are widely used as independent techniques to confirm PPIs. Both capture molecular complexes on solid-phase affinity matrices, followed by analyses using polyacrylamide gel electrophoresis (PAGE), western blotting and/or mass-spectrometry. Whilst these methodologies are currently considered gold-standards, they are biased toward the detection of high-affinity PPIs that remain associated with the affinity matrix throughout multiple, high-stringency washing steps (Gingras et al., 2007). Single Molecule Pulldown (SiMPull) can improve the detection of low affinity interactions by rapidly interrogating the affinity matrix with single molecule fluorescence nanoscopy to visualize transient binding events (Jain et al., 2011). A common drawback to all these approaches is that protein binding is assessed outside of the cell, under artificial buffer conditions and at non-physiological temperatures. Since buffer conditions alone can be a significant cause of false-negatives (Hakhverdyan et al., 2015), it is beneficial to assess candidate interactions in live cells. This allows interactions to be tested under native buffer and temperature conditions, taking advantage of intramolecular crowding effects and maintaining cytosolic concentrations of protein, which may physiologically help promote the formation of low-affinity complexes.

The use of proximity ligase techniques (BioID) or the combination of IP/AP with chemical crosslinking *in vivo* can be used to identify PPIs in live cells (Roux et al., 2012). PPIs can also be detected within live cells using microscopy-based techniques, including fluorescence correlation spectroscopy (FCS), fluorescence resonance energy transfer (FRET) and bimolecular fluorescence complementation (BiFC) (Kerppola, 2006, Ries et al., 2012, Wallrabe et al., 2005). Whilst these techniques provide detailed kinetic and structural information about the interaction, they require a combination of specialized imaging equipment, detailed optimization of fluorescent probe placement, and/or complex data post-processing. The fluorescence 2-hybrid approach (F2H/F3H) is a simpler assay that targets a fluorescently tagged query protein to a genomic DNA scaffold within the cell nucleus, and examines whether a query binding partner is recruited (Herce et al., 2013, Zolghadr et al., 2008). The F2H system benefits from simple image acquisition, but requires proteins to be freely accessible to the nucleus.

We have developed a miniaturized version of the affinity pulldown that detects PPIs in live cells by harnessing the native intracellular trafficking machinery. Our assay, which we term nanoscale pulldown (NanoSPD), couples a fluorescent bait protein to a molecular motor that endogenously traffics along filopodia: elongated, actin-based membrane protrusions that extend beyond the periphery of eukaryotic cells (Mattila et al., 2008). We show that fluorescent interacting proteins are forced to move synchronously along actin filaments in filopodia shafts, and that as a result of continuous co-transport, entire macromolecular complexes are actively concentrated at the distal tips of filopodia. The active concentration into a well-defined nanoscopic volume amplifies the fluorescence signal and allows for a simple quantitative readout using standard epifluorescence or confocal microscopy. We use nanoscale pulldowns to explore the interactome associated with taperin (TPRN), a protein required for detecting sound and mutated in a form of human hereditary hearing loss (Li et al., 2010, Rehman et al., 2010). Our data reveal a molecular link with CHARGE syndrome and indicates the existence of a potential nuclear-stereocilia signaling axis in mechanosensory hair cells.

## RESULTS

### Harnessing myosin-powered intracellular traffic to artificially deliver molecules within filopodia

Myosins are a superfamily of ATPase molecular motors that generate force on actin filaments to power a variety of biological processes, including cytokinesis, cell motility and intracellular trafficking (Sellers, 2000, Sweeney et al., 2010). Myosin motors are precisely targeted with the cell (Hartman et al., 2011), with sub-classes III, X and XV being specialized for delivering molecular cargos within filopodia and/or mechanosensory hair cell stereocilia (Belyantseva et al., 2003, Belyantseva et al., 2005, Berg et al., 2002, Fang et al., 2015, Les Erickson et al., 2003, Manor et al., 2011, Salles et al., 2009, Schneider et al., 2006, Tokuo et al., 2004, Zhang et al., 2004). We hypothesized that this normal physiological process of intracellular trafficking could be exploited to artificially target bait proteins within the cell, and then examine whether candidate prey proteins were also recruited.

This idea was explored using the class X myosin molecular motor (MYO10), which we chose due to its particularly robust ability to traffic along filopodia actin filaments (Berg & Cheney, 2002, Berg et al., 2000, Bohil et al., 2006). The MYO10 molecule consists of a core ATPase ‘motor’ that binds to actin filaments, a neck domain that associates with regulatory light chains, a coiled-coil motif that may promote dimerization and a C-terminal tail domain that provides the molecular scaffolding for cargo protein binding (Kerber et al., 2011) (Figure 1A). Similar to full-length MY010, truncated molecules (referred to as heavy meromyosin, MY010^hmm^, Figure 1A) consisting of the motor domain, IQs, and the coiled-coil still traffic and accumulate at filopodia tips (Berg & Cheney, 2002) (Figure 1B). The C-terminal tail domain is therefore not required for MY010 to traffic within filopodia and target to its tips. Next, we investigated whether replacing the endogenous cargo binding tail domain of MY010 with a ‘bait’ protein could similarly allow chimeric MY010-bait molecules to traffic along filopodia, potentially in complex with interacting ‘prey’ proteins.

**Figure 1.**
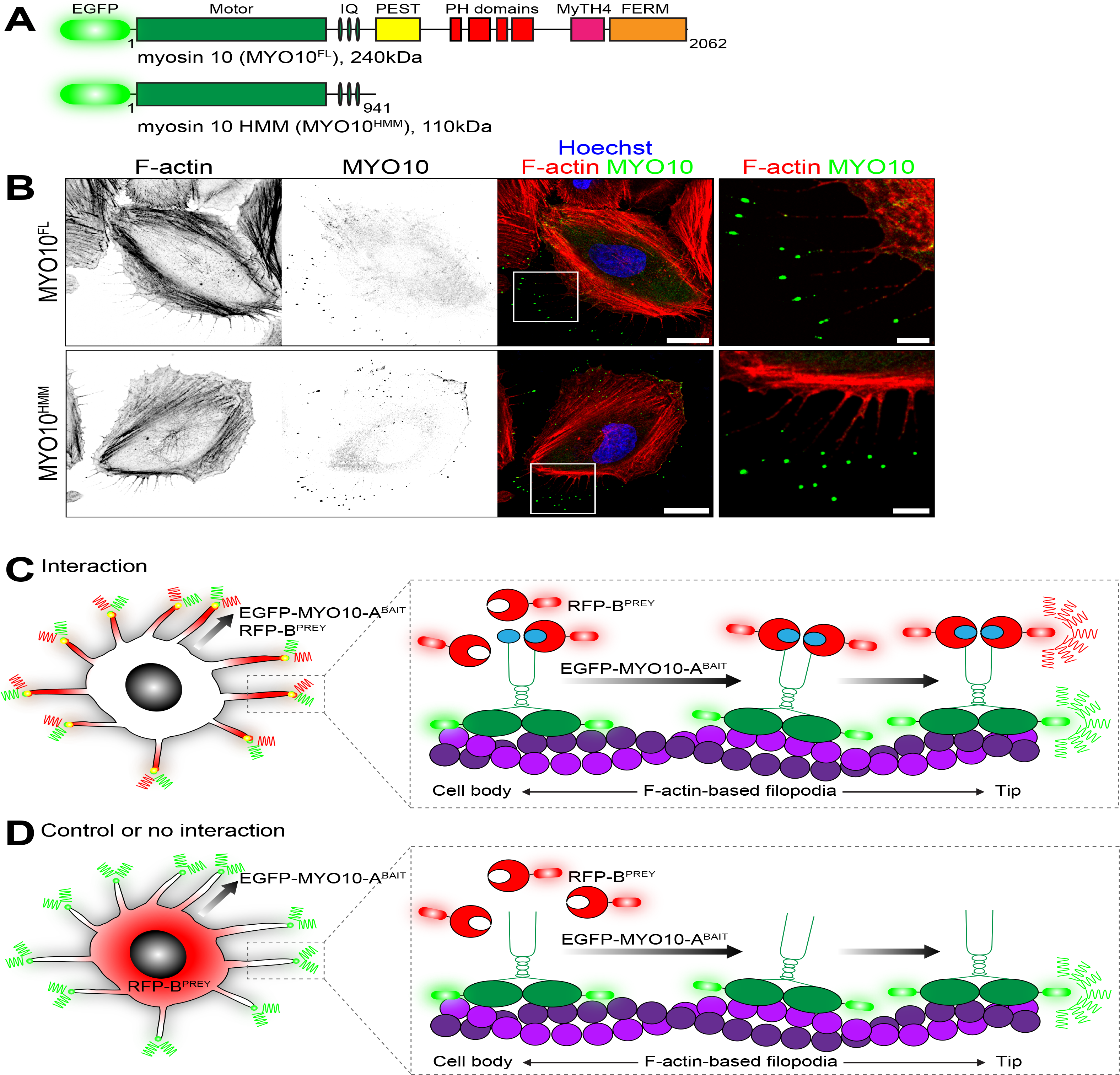
**Principle of the nanoscale pulldown (A)** Domain representation of full-length EGFP-tagged MY010^FL^ with the core ATPase motor domain, IQ motifs that bind light chains, a coiled-coil and tail domains are highlighted. Heavy meromyosin MY010^HMM^ molecules are identical, but are truncated after the coiled-coil. Nanoscale pulldown baits are constructed by fusing proteins in frame to the C-terminus of MY010^HMM^, effectively providing an alternate tail domain. **(B)** EGFP-tagged MY010^FL^ (top panel) and truncated MY010^HMM^ (bottom panel) both traffic to concentrate at the tips of filopodia in HeLa cells. Inset white boxes are magnified to show individual filopodia. F-actin is labeled with rhodamine phalloidin (red) and nuclei are stained with Hoechst (blue). Scale bar, 20 μm, 5 μm in magnifications. F-actin and MY010 images have been inverted. **(C)** Using nanoscale pulldown to test the interaction between proteins A^BAIT^ and B^PREY^. In this example, A^BAIT^ is forcibly trafficked to filopodia tips by MY010, as part of the EGFP-MY010-A^BAIT^ fusion protein (green). In the presence of an interaction, RFP-B^PREY^ molecules (red) are actively transported along filopodia, resulting in concentration of bait and prey molecules at filopodia tips. **(D)** In the control, EGFP-MY010^NO BAIT^ molecules (green) lacking a bait protein still accumulate at filopodia tips, but can no longer traffic RFP-B^PREY^ (red). Identical results are expected if RFP-B^PREY^ does not interact with EGFP-MY010-A^BAIT^.

For proof of principle, we tested the known interaction between the tail domain of MYO7A and the MYO7A and Rab interacting protein (MYRIP) (El-Amraoui et al., 2002, Fukuda et al., 2002). Since filopodia trafficking of wild-type MY010 occurs in the single molecule regime (Kerber et al., 2009), we used time-lapse total internal reflection fluorescence microscopy (TIRFM) to try and capture these events. Two classes of behavior were observed in HeLa cells expressing a chimeric EGFP-MY010^HMM^ bait fused to the tail domain of MYO7A, referred to as MY010-MY07A(TAIL)^bait^, and the prey mCherry-MYRIP, referred to as MYRIP^prey^. First, EGFP and mCherry signals specifically co-accumulated at the tips of filopodia. Second, we observed correlated bidirectional motility of EGFP and mCherry positive puncta along the filopodia shaft (Figure 2A). Bright fluorescent puncta moved slowly towards the cell soma (retrogradely), whilst very dim puncta moved anterogradely toward the filopodia tip (Figure 2B). These trafficking events were qualitatively similar to wild-type MY010, where retrograde motility is linked to treadmilling of the underlying actin filaments and anterograde motion results from active trafficking by the myosin motor (Kerber & Cheney, 2011, Kerber et al., 2009). As a control, we time-lapse imaged HeLa cells expressing MYRIP^PREY^ and EGFP-MY010, lacking the MYO7A-tail moiety (referred to as MY010^NO BAIT^) (Figure 2C). Similar to before, EGFP-positive MY010^NO BAIT^ puncta still moved bidirectionally along the shaft and strongly accumulated at filopodia tips. However, mCherry-positive puncta no longer trafficked along the filopodia shaft or accumulated at the tips (Figure 2D). We conclude that the MY010-MY07A(TAIL)^BAIT^ bait chimera actively co-transported MYRIP^PREY^ within filopodia and that correlated motility of bait-prey complexes can be used as a specific measure of protein-protein binding. We call this method nanoscale pulldown (or NanoSPD), since bait-prey complexes are forced to traffic, or are ‘pulled’, along filopodia by the MY010 molecular motor.

**Figure 2.**
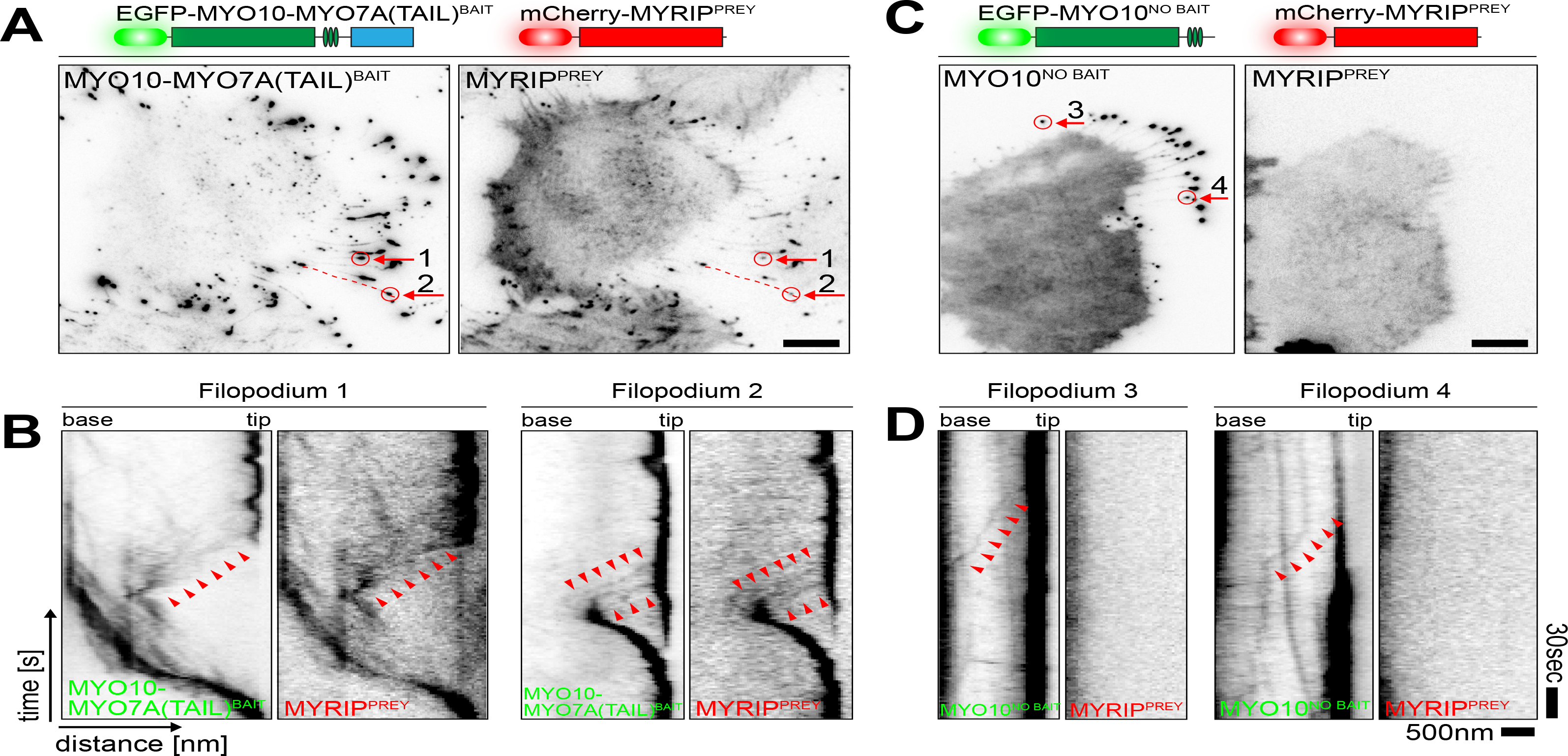
**Principle of the nanoscale pulldown** Visualization of bait and prey molecule trafficking in live HeLa cells by time-lapse total internal reflection fluorescence microscopy (TIRFM). Imaging was performed at 37°C. **(A)** In HeLa cells expressing MY010-MY07A(TAIL)^BAIT^ (left panel) and MYRIP^PREY^ (right panel), both molecules simultaneously accumulate at filopodia tips. A single time point from the time-lapse data set is shown. Filopodia selected for kymographs are indicated by red arrows. **(B)** Kymographs (plotting fluorescence intensity along a single filopodium vs. time) from (A) reveal molecular trafficking events. Bait and prey molecules accumulate at the filopodia tips which are visible as a dark vertical line. Very dim puncta of MY010-MY07A(TAIL)^BAIT^ and MYRIP^PREY^ move synchronously towards the filopodia tip (anterograde motion, arrow heads). Retrograde flow of bait and prey molecules (towards the cell soma) is also observed. **(C)** MY010^NO BAIT^ accumulates robustly at filopodia tips but is unable to traffic MYRIP^PREY^ molecules that remain diffuse in the cytoplasm. This control experiment demonstrates that trafficking of MYRIP^PREY^ is critically dependent on the MY07A(TAIL) moiety being fused to the myosin motor. **(D)** Kymographs from (C) confirm that MYRIP^PREY^ molecules are no longer synchronously trafficked by MY010^NO BAIT^. Anterograde (arrowheads) and retrograde (not shown) trafficking of MY010^NO BAIT^ is still observed. Filopodia 3 and 4 are indicated by arrows in (C). All fluorescence images have been inverted. Horizontal scale bars are 10 μm (A,C) and 500 nm (B,D), vertical scale bars are 30 seconds (C,D).

### Principle of the nanoscale pulldown

To develop a simple and sensitive readout for the nanoscale pulldown, we focused on quantifying the fluorescence intensity (i.e. the concentration) of bait-prey complexes at filopodia tips. This location proved advantageous as the sustained anterograde MY010 motor flux provided active concentration of bait-prey complexes, effectively working as a biological amplifier to boost fluorescence intensity. This unique amplification feature of nanoscale pulldowns allows filopodia to be imaged using standard confocal microscopy, rather than using more complex TIRFM in live cells. To test the interaction between two proteins A^BAIT^ and B^PREY^ by nanoscale pulldown, cells are transfected with a MY010-A^BAIT^ fusion protein and B^prey^, with the expectation that both molecules will accumulate at the filopodia tips (Figure 1C). A single control is required to confirm that B^PREY^ trafficking is dependent upon the specific interaction with the myosin-powered A^bait^. This is assessed by transfecting cells with MY010^HMM^ (lacking the bait domain and referred to as MY010^NO BAIT^) and B^PREY^ (Figure 1D). All molecules used in this study are tagged with spectrally orthogonal fluorescent proteins and are not explicitly stated in the main text (see Methods and Figures).

The nanoscale pulldown methodology was initially tested in HeLa cells using the MYO7A – MYRIP interaction as a proof of principle. HeLa cells expressing MYRIP^prey^ and either MY010-MY07A^bait^ or MY010^no BAIT^ were fixed and imaged using confocal microscopy (Figure 3A). Analogous to our observation in live cells (Figure 2), MYRIP^prey^ was robustly transported to filopodia tips by MY010-MY07A^bait^, but not by MY010^no BAIT^ (Figure 3A). Similar observations were made in COS-7 cells (data not shown) and also in Sf9 (*Spodoptera frugiperda*) insect cells (Figure 3D), demonstrating that nanoscale pulldowns may be broadly applicable to a range of mammalian and non-mammalian cell lines. Whilst the assay worked equivalently in all cell lines tested, Sf9 insect cells were particularly notable for reproducibly generating large numbers of filopodia per cell.

**Figure 3.**
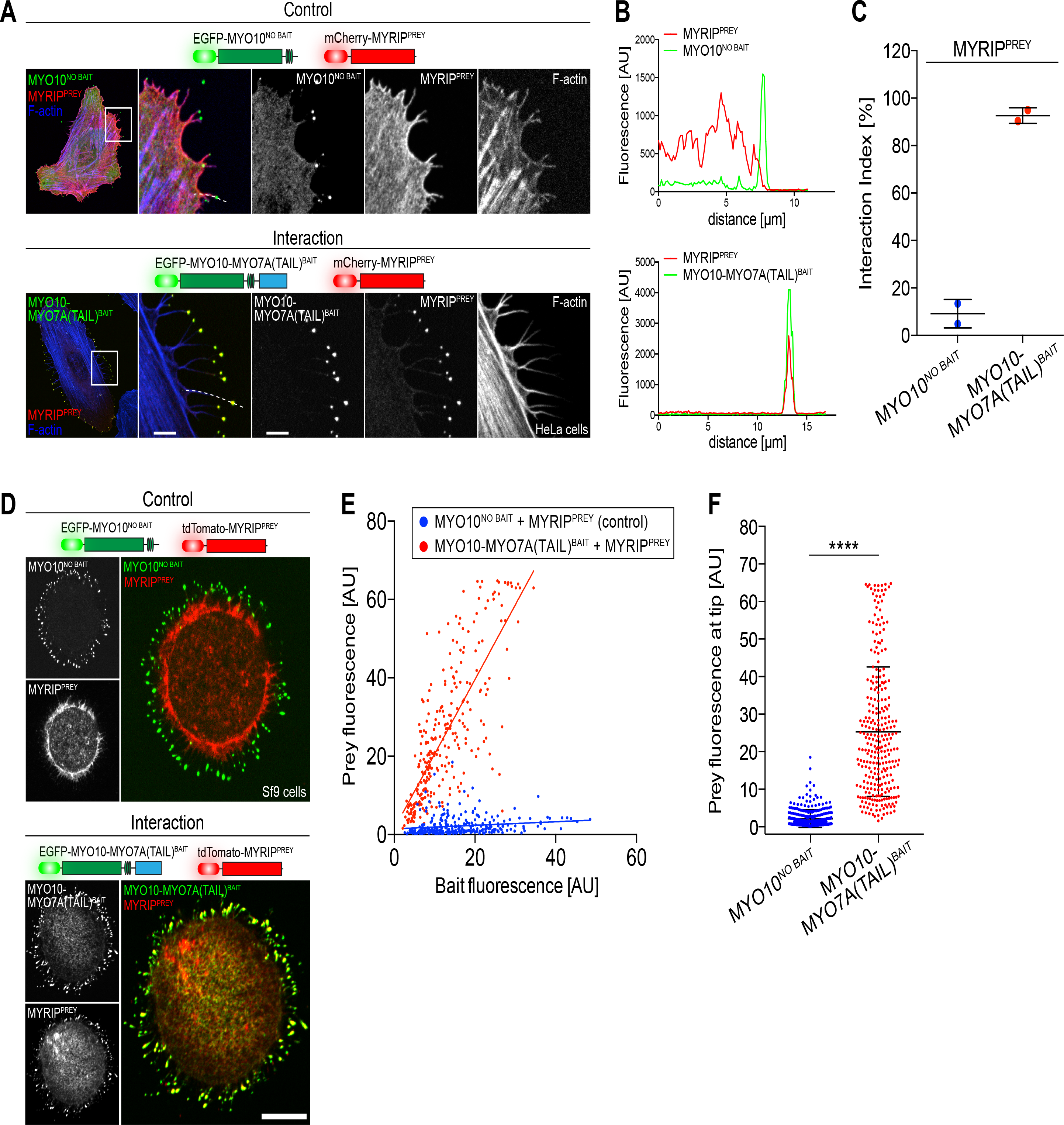
**Quantifying PPIs by nanoscale pulldown** Continuous trafficking by MY010 concentrates bait-prey complexes into a nanoscopic volume at the filopodia tips. This unique feature of the nanoscale pulldown yields real biological (noise-free) signal amplification and provides a defined location for optically interrogating a molecular interaction with minimal background fluorescence. **(A)** In chemically fixed HeLa cells expressing MY010-MY07A(TAIL)^BAIT^ and MYRIP^prey^, both bait and prey molecules co-accumulate at filopodia tips. MY010^NO BAIT^ is unable to traffic MYRIP^PREY^ to the tips in the control, confirming the specificity of the interaction and that MYRIP^PREY^ cannot traffic autonomously. Scale bars, 20 μm; magnified boxed regions, 5 μm. **(B)** Using spatial correlation to detect molecular interactions. A line scan along the filopodia shaft (dotted line) is used to calculate Pearson’s *r* correlation coefficient between the bait and prey fluorescence signals. The likelihood that the observed correlation could have occurred by random chance is estimated by bootstrapping (see Materials and Methods). **(C)** The interaction index (# significantly correlated filopodia / total # filopodia) is calculated for each experimental condition. Co-expression of MYRIP^PREY^ with MY010-MY07A(TAIL)^BAIT^ yields a significantly higher interaction index than when MYRIP^PREY^ is co-expressed with MY010^NO BAIT^. Each data point is the average interaction index from a single experimental determination. **(D)** Nanoscale pulldowns can be used in diverse cell types, including Sf9 insect cells that produce large numbers of filopodia. In fixed Sf9 cells, MYRIP^PREY^ accumulates with MY010-MY07A(TAIL)^BAIT^ at the filopodia tips, but not when co-expressed with MY010^NO BAIT^, similar to HeLa cells in (A). Scale bar, 10 μm. **(E)** Using intensity correlation to detect molecular interactions in Sf9 cells. Scatter plot of bait (x-axis) and prey (y-axis) fluorescence at individual filopodia tips measured from line scans. MYRIP^prey^ fluorescence at the filopodia tips is strongly correlated with MY010-MY07A(TAIL)^bait^ fluorescence (Pearson’s *r* = 0.77), unlike cells expressing MY010^NO BAIT^ (*r* = 0.16). **(F)** Bar-graph plot of MYRIP^prey^ fluorescence intensities from data in (D). Co-expression with MY010-MY07A(TAIL)^bait^ significantly increases the average fluorescence at filopodia tips, compared to MY010^NO BAIT^ (Mann-Whitney *U* test).

### Quantification of nanoscale pulldowns to test protein-protein interactions

We developed a quantitative framework and software tool to objectively test for spatial correlation between bait and prey molecules using Pearson’s correlation coefficient (*r*). Line scans along individual filopodia were used to measure the fluorescence intensity of bait and prey molecules (Figure 3B). In the presence of an interaction, bait and prey fluorescence intensities are expected to be correlated (i.e. *r* = +1), whereas these quantities should be uncorrelated in the absence of an interaction (i.e. *r* = 0). However, calculated *r*-values for interacting bait-prey pairs were frequently intermediate between 0 and 1 (data not shown). Using an arbitrary global threshold to assign significance to these intermediate values was unsatisfactory, since partial correlations may still indicate a significant biological interaction. Instead, we estimated the likelihood that any measured correlation occurred through chance by using bootstrapping to randomize the prey signal and calculate the exact distribution of *r*-values for all possible permutations (Costes et al., 2004, Lifshitz, 1998). This approach allowed us to test the null hypothesis that a given bait-prey correlation in an individual filopodium occurred randomly. Detailed documentation of this algorithm, MATLAB source code and user instructions are available in the Supplementary Materials.

The Pearson’s *r*-based quantification algorithm was tested on line scans from HeLa cells expressing MYRIP^prey^ and either MY010-MY07A^bait^ or MY010^no BAIT^ (Figure 3B). In cells expressing MY010-MY07A^bait^ and MYRIP^prey^ we found that 93 ± 3% of filopodia (192 filopodia sampled, two independent trials) had significantly correlated bait-prey fluorescence intensities (Figure 3C). Conversely, in control HeLa cells expressing MY010^no BAIT^ and MYRIP^PREY^, only 9 ± 6% of filopodia (164 filopodia sampled, two independent trials) were significantly correlated. The ~10-fold increase in interaction index (percentage of total filopodia with bait-prey correlation) was statistically significant (*p* = 0.01, 2-tailed *t*-test), and quantitatively confirmed the interaction between these two proteins (Figure 3C).

While the Pearson’s quantification algorithm was sensitive to spatial correlations of bait-prey complexes in HeLa cells, it reported a higher level of false-positive correlations in Sf9 cells when used to evaluate control interactions such as MY010^no BAIT^ and MYRIP^prey^ (data not shown). The likely cause of this phenomenon was from cytoplasmic volume filling of the filopodia tips, which were shorter and more bulbous in Sf9 cells. Since Pearson’s *r* coefficient does not consider the magnitude of fluorescence changes, small increases in prey fluorescence due to volume-filling are detected as an artefactual correlation. To address this issue in Sf9 cells, we measured the absolute fluorescence intensities at filopodia tips. When examined over a large sample of independent filopodia, bait and prey fluorescence intensities are expected to be correlated in the presence of an interaction, and uncorrelated otherwise. A critical condition of this intensity-based analysis is that imaging conditions are standardized (objective, emission filters, detector gain/exposure, laser power) so that data from independent experiments can be combined.

Line scan data from Sf9 cells testing the MYO7A – MYRIP interaction (Figure 3D) were reanalyzed using the intensity-based correlation algorithm. A linear correlation between bait and prey fluorescence intensity was observed at the filopodia tips of cells expressing MY010-MYO7A^BAIT^ and MYRIP^prey^ (red) (Figure 3E). By comparison, in cells expressing MY010^no bait^ and MYRIP^prey^, bait and prey fluorescence values were uncorrelated (blue) (Figure 3E). In addition to correlation, the average fluorescence intensity of MYRIP^prey^ at filopodia tips was increased by 11.8-fold in the presence of MY010-MY07A^bait^ compared to MY010^no bait^ (Figure 3F) (*p* < 0.0001, Mann-Whitney *U* test), providing a separate measure of the interaction by nanoscale pulldown. We considered the possibility that the difference in MYRIP^prey^ fluorescence at tips could be due to differences of trafficking efficiencies between MY010-MY07A^bait^ and MY010^no BAIT^. While there was a statistically significant difference between the mean fluorescence intensities of MY010^no BAIT^ versus MY010-MY07A^bait^, the effect size (1.2-fold) was minimal (Supplementary Figure S1A). The large increase in MYRIP^prey^ fluorescence was therefore due to the specific interaction with MY010-MYO7A^BAIT^. We conclude that the intensities of bait and prey fluorescence within filopodia can also be used as a sensitive measure of a protein-protein interaction in nanoscale pulldowns.

### Nanoscale pulldowns can traffic multiple preys to dissect binding hierarchies in macromolecular complexes

We next investigated if nanoscale pulldowns could be expanded to traffic multiple prey species along filopodia and thus probe the composition of more complex macromolecular assemblies. We examined the tripartite complex between the C-terminal tail domain of myosin 5a (MYO5A) and the Ras-related GTPase RAB27A, that requires the effector melanophilin (MLPH) as an intermediate adaptor (Wu et al., 2002) (Figure 4A). This interaction was tested by nanoscale pulldown in Sf9 cells. Qualitatively, RAB27A^PREY^ did not concentrate at filopodia tips in Sf9 cells expressing either MY010-MY05A(TAIL)^BAIT^ (Figure 4B) or MY010^NO BAIT^ (data not shown). This confirmed that RAB27A^PREY^ failed to traffic in the absence of the critical adapter MLPH. When a non-fluorescent (dark) myc-tagged MLPH molecule was introduced, RAB27A^PREY^ was trafficked to filopodia tips by MY010-MY05A(TAIL)^bait^, confirming that MLPH was critical for assembly of the MYO5A melanosome receptor (Figure 4B). Data were analyzed using the intensity-based correlation algorithm. The average fluorescence intensity of MY010-MY05A(TAIL)^BAIT^ molecules at filopodia tips was independent of the presence, or absence of myc-MLPH, indicating no systematic change in bait molecule trafficking (Supplemental Figure S1B). Despite the equal trafficking of MY010-MY05A(TAIL)^BAIT^ molecules, there was a significant 6.5-fold increase in RAB27A^PREY^ fluorescence in the presence of myc-MLPH (Figure 4C). We conclude that the association of RAB27A with MYO5A(TAIL) was critically dependent upon the effector MLPH. Our data show that a myosin-powered bait can traffic multiple, interacting prey molecules within filopodia, and that this can be exploited by nanoscale pulldowns to study the composition and hierarchy of larger macromolecular complexes.

**Figure 4.**
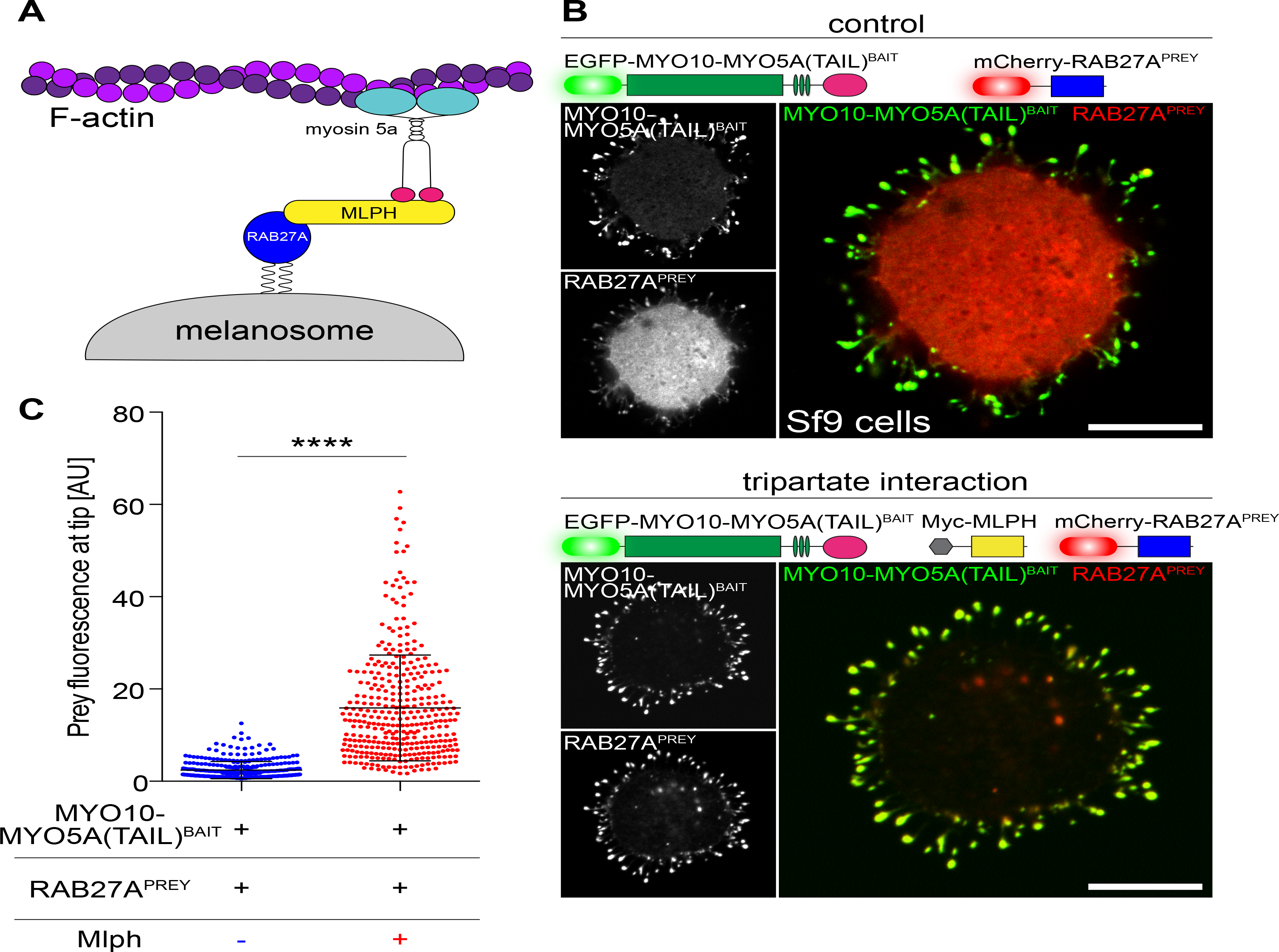
**Dissecting hierarchies in macromolecular complexes by nanoscale pulldown (A)** Schematic of the tripartite melanosome receptor formed by MY05A, MPLH and RAB27A. Active RAB27A (blue) attaches to the melanosome membrane via geranylgeranyl moieties, binds its effector melanophilin (MLPH, yellow), which is then recruited to the MY05A tail domain (pink). This tripartite complex is trafficked along actin filaments by the MY05A motor domain (cyan). **(B)** Using nanoscale pulldown in Sf9 cells to test the requirement for MLPH in complex formation. RAB27A^PREY^ is not robustly trafficked by MY010-MY05A(TAIL)^bait^, despite the bait accumulating strongly at filopodia tips (top panel). Inclusion of myc-tagged MLPH (dark) results in co-accumulation of both MY010-MY05A(TAIL)^BAIT^ and RAB27A^PREY^ at filopodia tips, demonstrating MLPH is required for complex formation (bottom panel). Scale bars, 10 μm. **(C)** Intensity correlation analysis of RAB27A^prey^ fluorescence at filopodia tips from experiments shown in (B). RAB27A^prey^ tip fluorescence is significantly increased upon coexpression of Myc-MLPH (Mann-Whitney *U* test), confirming the interaction.

### Using nanoscale pulldowns to explore the taperin interactome and molecular mechanisms of hearing loss

Having experimentally validated the nanoscale pulldown technique, we used it to study protein-protein interactions and macromolecular complexes implicated in hearing and pathophysiology that causes deafness. Genetic studies of human hereditary hearing loss in conjunction with proteomic analyses have identified an extensive catalog of proteins involved in the detection of sound by the cochlea (Barr-Gillespie, 2015, Richardson et al., 2011). Many of these proteins assembly into macromolecular complexes within auditory hair cell stereocilia, mechanosensory organelles that are the primary transducers of sound (Schwander et al., 2010). We previously reported that taperin (TPRN) concentrates within stereocilia and is mutated in an inherited form of autosomal recessive human deafness (Rehman et al., 2010). The molecular function of taperin remains unknown in hair cells, although its localization suggests that it might be a component of ankle-link complexes, large macromolecular assemblies that interconnect neighboring stereocilia at their base (Rehman et al., 2010, Salles et al., 2014).

To gain insight into taperin’s function in mechanosensitive hair cells, we screened Y2H libraries prepared from mouse inner ear and kidney RNA using full-length taperin as bait (Supplemental Table S2). We studied a subset of these interactors in more detail, not only to confirm their validity by orthogonal technique, but also to test the nanoscale pulldown assay with a variety of different preys, such as membrane-associated and nuclear proteins. The Y2H assays identified the alpha (PPP1CA) and gamma (PPP1CC) catalytic subunits of protein phosphatase 1 (PP1) as taperin interacting proteins. PP1 is a widely expressed serine/threonine phosphatase and regulates a variety of cellular functions, including glycogen metabolism, transcription, protein synthesis, cell division, and apoptosis. PP1 is a multimeric enzyme consisting of a highly conserved catalytic and several variable regulatory units. PP1 isoforms and its subunits can be located throughout the cell, including in the nucleus (Rebelo et al., 2015). We tested the interaction of the catalytic subunit of PP1 and taperin by nanoscale pulldown in Sf9 cells expressing MY010-TPRN^BAIT^ and either PPP1CA^PREY^ (alpha), PPP1CB^PREY^ (beta) or PPP1CC^PREY^ (gamma) subunits (Figure 5A,B + S2B,C). Intensity-based analysis revealed that the mean fluorescence intensities of PPP1CA^prey^, PPP1CB^PREY^ and PPP1CC^PREY^ at filopodia tips were significantly increased by 5.2-fold, 1.3-fold and 5.7-fold, respectively, when expressed with MY010-TPRN^BAIT^ versus MY010^NO BAIT^ (Figure 5C). Although statistically significant, the fluorescence increase (i.e. the effect size) of PPP1CB was minimal (1.3-fold) and indicated a preference of taperin to bind PPP1CA and PPP1CC. These observations mirrored the identification of PPP1CA and PPP1CC by Y2H (Supplemental Table S2) and confirm a previous report (Ferrar et al., 2012). We conclude that nanoscale pulldowns can successfully detect variations in binding affinities between members of a conserved protein family.

**Figure 5.**
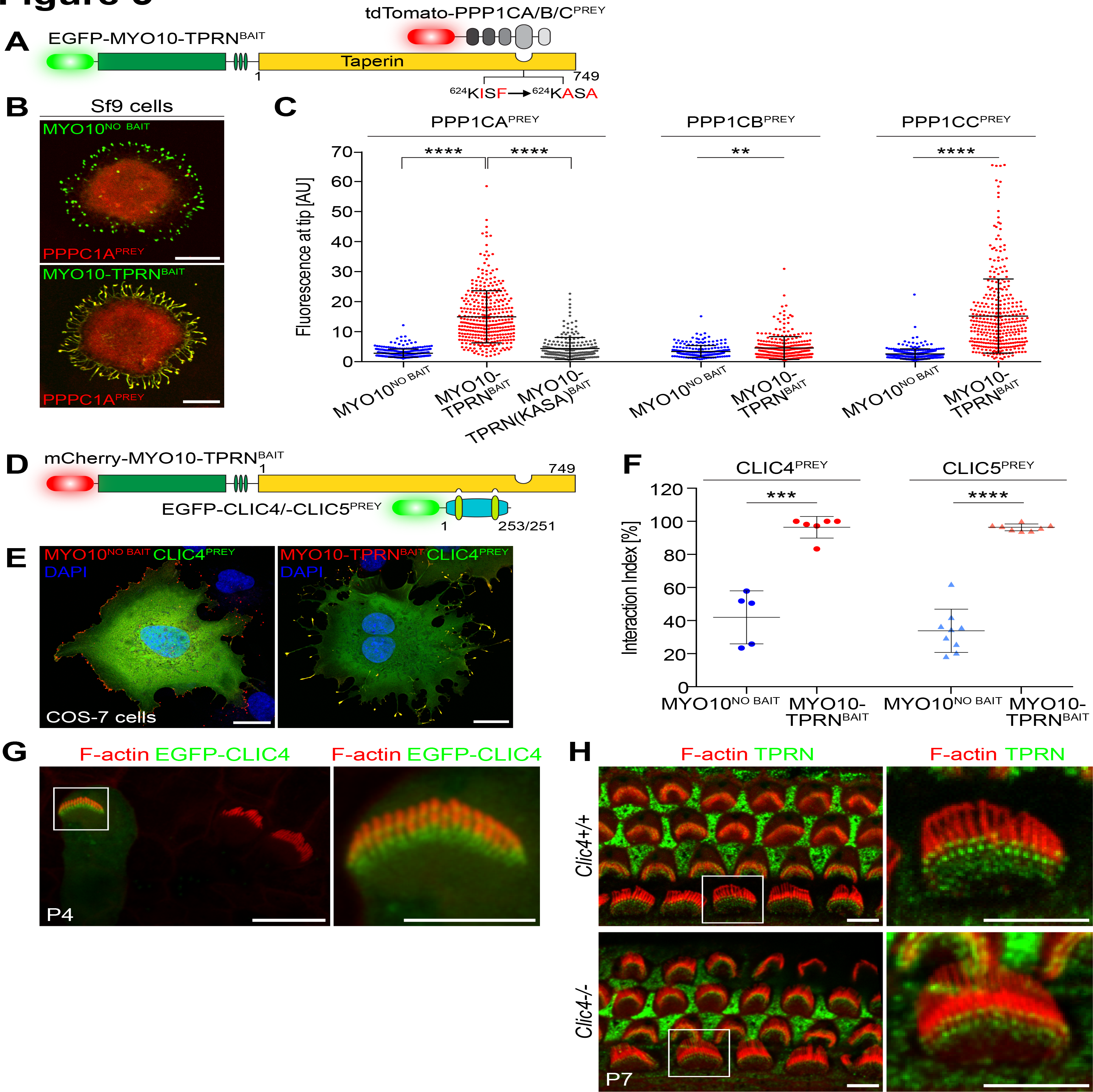
**Using nanoscale pulldowns and mutagenesis to study taperin binding to PP1 and CLIC proteins (A)** Molecules used to test interactions between taperin (TPRN) and the catalytic subunits of protein phosphatase 1 (PPP1CA, PPP1CB, PPP1CC) by nanoscale pulldown. Point mutations in taperin (KISF → KASA) that disrupt the conserved PP1 binding motif are indicated. **(B)** MY010-TPRN^BAIT^ delivers PPP1CA^PREY^ to the filopodia tips in Sf9 cells (bottom panel). MY010^NO BAIT^ is unable to traffic PPP1CA^PREY^ despite accumulating robustly at the filopodia tips (top panel), confirming the specificity of the assay. **(C)** Intensity correlation analysis reveals selective binding of taperin to PP1 catalytic subunits. Strong accumulation of PPP1CA^prey^ and PPP1CC^prey^ was detected at the filopodia tips of Sf9 cells expressing MY010-TPRN^BAIT^, as compared to MY010^NO BAIT^ (PPP1CA, ANOVA; PPP1CB/PPP1CC, Mann-Whitney *U* test). Note: although the increase of PPP1CB^prey^ fluorescence was statistically significant, the effect size was minimal, indicating no substantial interaction. Nanoscale pulldowns were also used to test the effect of mutating the binding interface for PPP1CA within taperin (KISF → KASA). The fluorescence intensity of PPP1CA^prey^ was significantly reduced in cells expressing MY010-TPRN(KASA)^BAIT^, compared to MY010-TPRN^BAIT^ (ANOVA). Data are quantified from panel (B) and Figure S2. **(D)** Nanoscale pulldown molecules to test the interaction of taperin with CLIC family proteins. For these experiments, bait constructs are tagged with mCherry, and prey constructs are tagged with EGFP. **(E)** CLIC4^PREY^ is trafficked to filopodia tips by MY010-TPRN^BAIT^ (right panel) in mammalian COS-7 cells, but not by MY010^NO BAIT^ (left panel). Similar observations were made for CLIC5 (see Figure S3). **(F)** Quantification of the taperin interaction with CLIC4/5 using fluorescence spatial correlation. Statistically significant increases in the interaction index for CLIC4 and CLIC5 were observed when MY010-TPRN^BAIT^ was expressed compared to MY010^NO BAIT^ (*t*-test). Each data point represents one experimental determination (n = 267 – 516 filopodia total). **(G)** Biolistic expression of EGFP-CLIC4 (green) in cultured explants of P4 cochlea. CLIC4 localizes towards the base of actin-based stereocilia where they insert into the apical pole of the mechanosensory hair cell. Image shown is 3D orthogonal reconstruction of confocal z-stack. F-actin is labeled with rhodamine phalloidin (red). White box indicates transfected hair cell and is shown magnified (right panel). **(H)** Immunofluorescence detection of taperin (green) in the mouse cochlea at P7. In wild-type cochleae (top row), taperin is detected at the base of hair cell stereocilia (boxed inset and magnified), and in supporting cells. Despite interacting with CLIC4, taperin localization is indistinguishable in *Clic4^-/-^* null littermates (bottom row). F-actin is labeled with rhodamine phalloidin (red). Scale bars, 10 μm (B, G); 20 μm (E); 5 μm (H, magnified image in G).

### Nanoscale pulldown can detect changes in binding affinity due to point mutations

Taperin contains a consensus KISF motif (residues 624 – 627) that binds to a hydrophobic patch within the catalytic domain of PP1 isozymes to inhibit phosphatase activity (Ferrar et al., 2012). Mutation of critical residues in this KISF motif to alanine (KISF to KASA) abolishes taperin binding to PPP1CA when measured by conventional affinity-pulldowns (Ferrar et al., 2012). We examined whether nanoscale pulldowns were sensitive enough to detect a change in binding caused by targeted point mutations. Intensity-based analysis in Sf9 cells showed a statistically significant 3.3-fold reduction in PPP1CA^PREY^ accumulation at filopodia tips when co-expressed with MY010-TPRN(KASA)^BAIT^ versus wild-type MY010-TPRN^BAIT^ (Figure 5C + S2A). Although MY010-TPRN(KASA)^BAIT^ significantly reduced the accumulation of PPP1CA^prey^, we still detected a small 1.6-fold increase in PPP1CA^PREY^ accumulation in Sf9 cells co-expressing MY010-TPRN(KASA)^BAIT^ versus MY010^NO BAIT^. These data indicate that the KASA mutation lowered the affinity of taperin binding to PPP1CA, but did not inhibit the interaction completely. We conclude that nanoscale pulldowns can detect the functional effect of critical amino acid substitutions in a conserved binding motif.

### Testing membrane-associated interactions by nanoscale pulldown

The Y2H screen also identified chloride intracellular channels 1, 4 and 5 (CLIC1, CLIC4, CLIC5) as potential taperin interacting proteins (Supplementary Table S2). The CLIC protein family members are highly conserved in vertebrates and exists as both soluble globular proteins and integral membrane channels. As integral membrane proteins, CLIC proteins may act as chloride channels (Berryman et al., 2004, Singh et al., 2007), while their function as globular soluble proteins is not well understood (Littler et al., 2010). CLICs have been shown to regulate diverse cellular processes such as receptor trafficking, tubulogenesis, angiogenesis, and apoptosis (Bohman et al., 2005, Maeda et al., 2008, Suh et al., 2004, Ulmasov et al., 2009). CLIC4 and CLIC5 are both highly expressed in inner ear hair cells and are detected in the stereocilia bundle proteome (Shen et al., 2015). Furthermore, immunofluorescence analysis showed that similar to taperin, CLIC5 protein is localized at the base of hair cell stereocilia (Gagnon et al., 2006). Consistent with our Y2H data, CLIC5 was reported to bind taperin by IP (Salles et al., 2014); however complex formation was critically dependent upon chemical cross-linking, suggesting that the interaction was normally low affinity and could dissociate from the affinity matrix.

We independently tested the binding of CLIC5 to taperin using nanoscale pulldown. COS-7 cells were transfected with CLIC5^PREY^ along with either MY010-TPRN^BAIT^ or MY010^NO BAIT^ (Figures 5D and S3) and analyzed using the Pearson’s *r*-based correlation algorithm. In cells expressing MY010-TPRN^BAIT^, 96.4% of filopodia exhibited significant correlation with CLIC5^prey^, but this was significantly reduced to 33.9% of filopodia when coexpressed with MY010^NO BAIT^ (Figure 5F). Using a similar experimental setup the interaction between CLIC4^PREY^ and MY010-TPRN^BAIT^ was confirmed (Figures 5D,E,F). Our data demonstrate that nanoscale pulldowns can successfully test binding with membrane-associated proteins, in addition to potentially revealing low affinity interactions in live cells that might otherwise escape detection in a conventional IP.

Given that CLIC4 is highly expressed in hair cells (Shen et al., 2015), we investigated whether CLIC4 protein was concentrated towards the base of stereocilia, similar to the distribution of taperin and CLIC5. In humans, the amino acid sequence of CLIC4 is 74.7% identical to that of CLIC5, making it difficult to develop isoform-specific antibodies. To specifically localize CLIC4, cochlear sensory epithelia were transfected with EGFP-tagged CLIC4 and imaged by confocal microscopy (Figure 5G). In transfected hair cells, EGFP-CLIC4 was observed concentrating at the base of stereocilia, consistent with it binding to taperin at this location. We hypothesized that CLIC4 might be involved in retaining, or stabilizing taperin within a larger macromolecular complex at the stereocilia taper region. We therefore examined the localization of taperin in both wild-type and *Clic4*^-/-^ null cochleae at postnatal day 7 (P7). Taperin localization was indistinguishable between wild-type and *Clic4*^-/-^ hair cells, indicating that CLIC4 was not required to establish taperin localization in stereocilia (Figure 5H). Consistent with the normal stereocilia architecture at P7, the hearing function of adult *Clic4*^-/-^ mutant mice was also grossly normal. Hearing sensitivities were measured at 4 weeks of age by auditory brainstem response (ABR) and showed that *Clic4*^-/-^ mice were indistinguishable from wild-type littermate controls (Supplemental Figure S4). We conclude that despite CLIC4 binding to taperin, and separately co-localizing with taperin at the base of stereocilia, the loss of this molecule from hair cells is not detrimental to auditory transduction. The presence of CLIC5 may be able to compensate for the loss of CLIC4 at the base of stereocilia in *Clic4*^-/-^ mice. Although we note that CLIC4 cannot reciprocally compensate for the loss of CLIC5 in the profoundly deaf *Clic5*^-/-^ *jitterbug* mutant mouse (Gagnon et al., 2006).

### Taperin binds chromodomain proteins to establish a potential nuclear-stereocilia signaling axis in auditory hair cells

Chromodomain helicase DNA binding protein 4 (CHD4) was the most abundant interaction identified by Y2H (Supplementary Table S2). CHD4 is a catalytic subunit of the nucleosome-remodeling and histone deacetylase (NuRD) complex that regulates nuclear DNA damage responses (Larsen et al., 2010, Polo et al., 2010). CHD4 belongs to a superfamily of nine CHD enzymes that contain tandem N-terminal chromodomains (<u>chr</u>omatin <u>or</u>ganization <u>mo</u>difier) and a central SNF2-like ATPase domain encompassing a DEAD-box like helicase superfamily domain (DEXDc) (Durr et al., 2006, Hall et al., 2007). We used nanoscale pulldowns in Sf9 cells to test the putative interaction between CHD4 and taperin identified by Y2H (Figure 6A). A CHD4 fragment (amino acids 561-936) identified in our Y2H screen was used to eliminate all but one of the predicted nuclear localization signals (NLS) to avoid potential accumulation of CHD4 in the nucleus. The fluorescence intensity of CHD4(561-936)^PREY^ at filopodia tips was 7-fold stronger in the presence of MY010-TPRN^BAIT^ versus MY010^NO bait^, quantitatively confirming the interaction (Figure 6B). The CHD4(561-936)^PREY^ fragment contains the second chromodomain and DEXDc domain. We further truncated this CHD4 fragment and found the DEXDc domain alone, CHD4(DEXDc)^PREY^, was sufficient to bind MY010-TPRN^BAIT^ and be transported to filopodia tips (Figure 6C). A reciprocal experiment was performed to isolate the region within taperin required to bind to the DEXDc domain of CHD4. Nanoscale pulldowns using CHD4(DEXDc)^PREY^ and a series of truncated MY010-TPRN^BAIT^ molecules encompassing residues p.1-260, p.261-622 or p.623-749 showed that the CHD4 DEXDc domain specifically interacted with the N-terminal 260 residues of taperin (Figure 6C).

We investigated whether taperin might be able to bind other CHD proteins, since the DEXDc domain is broadly conserved across superfamily members. Separate nanoscale pulldowns were performed in Sf9 cells using CHD1(DEXDc)^PREY^, CHD2(DEXDc)^PREY^, CHD3(DEXDc)^PREY^ or CHD7(DEXDc)^PREY^ in combination with either MY010-TPRN^BAIT^ or MY010^NO BAIT^ as control. Fluorescence intensity analyses at filopodia tips revealed that taperin bound to the DEXDc domains of CHD3 and CHD7, but not CHD1 and CHD2 (Figure 6D). We conclude that taperin can interact with multiple CHD proteins via the DEXDc domain, but exhibits selectivity towards specific protein family sub-members. More broadly, our results indicate that nanoscale pulldowns can be used to study nuclear proteins and to perform domain mapping experiments to understand structure/function relationships.

**Figure 6.**
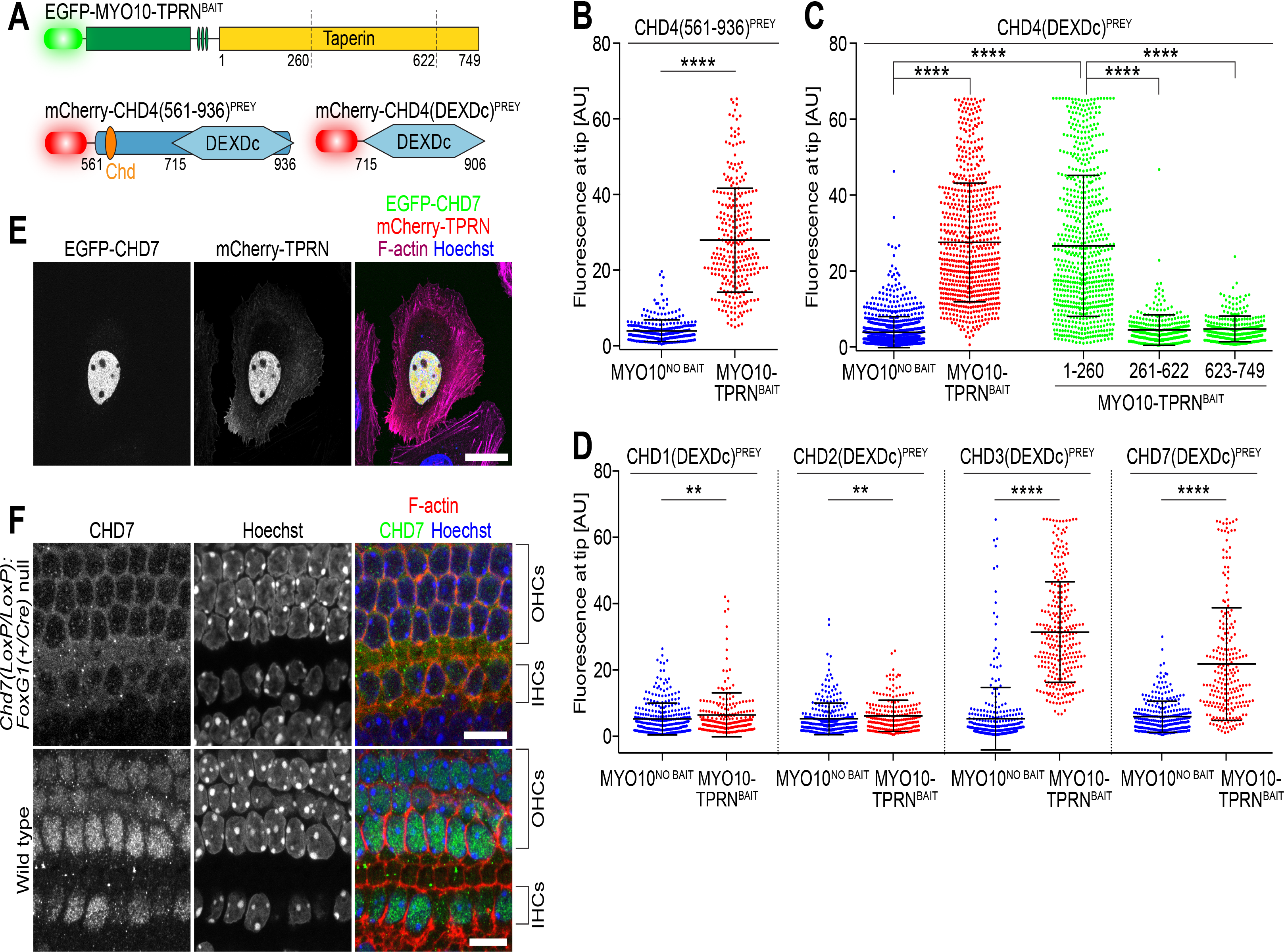
**Taperin interacts with chromodomain (CHD) family proteins in mechanosensory hair cells (A)** Nanoscale pulldowns molecules to test the interaction between taperin and CHD4. The CHD4 fragment (aa. 561 – 936) was the minimal domain identified by Y2H to interact with taperin. DEXDc = DEAD-like helicase domain; Chd = Chromodomain. Dotted lines demarcate truncation point for domain mapping experiments. **(B)** Intensity analysis in Sf9 cells confirms the statistically significant increase in CHD4(561-936)^prey^ fluorescence at filopodia tips when expressed with MY010-TPRN^BAIT^ (Mann Whitney *U*-test). **(C)** Domain mapping of the CHD4-TPRN interaction using nanoscale pulldowns. Intensity analysis in Sf9 cells show that the truncated CHD4(DEXDc)^PREY^ molecule containing the DEXDc helicase domain (aa. 715 – 906) alone was sufficient to bind taperin. Reciprocal truncations show that the N-terminal 260 amino acids of taperin were sufficient to bind and traffic the CHD4(DEXDc)^PREY^ domain. Other truncated regions of taperin (aa. 261-622 + aa. 623-749) were unable to traffic CHD4(DEXDc)^PREY^. Comparisons were made by one-way ANOVA with Tukey’s post-hoc test. **(D)** Taperin selectively binds to the DEXDc domain of CHD paralogs. Intensity analysis in Sf9 cells shows statistically significant binding of taperin to the DEXDc domains of CHD3, CHD4 and CHD7, but not to CHD1 or CHD2 (Mann-Whitney *U* test). **(E)** mCherry-taperin (red) and EGFP-CHD7 (green) co-localize to the nucleus when overexpressed in HeLa cells. F-actin is labeled with Alexa Fluor 633 phalloidin (magenta) and the nucleus is stained with Hoechst (blue). Scale bar, 25 μm. **(F)** Antibody labeling of CHD7 in P1 cochleae from wild-type and conditional null *Chd7^GT/LoxP^:FoxG1^Cre/+^* mice. OHCs = outer hair cell; IHCs = inner hair cells. Scale bars, 10 μm.

The interaction between taperin and CHD7 was intriguing since mutations in *CHD7* cause human CHARGE syndrome (OMIM: #214800), a pleiotropic childhood developmental disorder that includes deafness amongst multiple other pathological phenotypes (Martin, 2015). We hypothesized that CHARGE and DFNB79 non-syndromic deafness might share a similar etiology, potentially by affecting different components of a macromolecular complex critical for hearing. When expressed in HeLa cells, mCherry-TPRN and EGFP-CHD7 co-localized in the nucleus, indicating these binding partners may act on a common target or pathway (Figure 6E). To explore this hypothesis further we examined the distribution of CHD7 protein within the inner ear. CHD7 is expressed in neuro-epithelial cells of the developing cochlea at embryonic day 10.5 (E10.5), before hair cells have terminally differentiated and assembled mechanosensory stereocilia (Hurd et al., 2007, Hurd et al., 2010). We investigated the localization of CHD7 using immunofluorescence in postnatal day 1 (P1) mouse cochlea. CHD7 was prominently detected in the nucleus of mechanosensory hair cells at P1 (Figure 6F) and also at P14 (data not shown) indicating this was the stable distribution in functionally mature sensory cells. CHD7 antibody labeling was examined in cochleae from conditionally null *Chd7* mice carrying the *FoxG1^+/Cre^* allele to specifically knock-out expression of CHD7 in the inner ear and brain. Antibody labeling was absent from nuclei in conditional *Chd7* null cochleae, confirming the specific localization of CHD7 in sensory hair cells (Figure 6F).

Our data show that taperin can interact with CHD7 as part of the NuRD chromatin remodelling complex, and suggest that it may regulate gene transcription in mechanosensory hair cells. In support of this, we found that taperin interacts with PPP1CA (Figure 5)(Ferrar et al., 2012), which separately binds the IKAROS (IKZF1) zinc-finger protein to regulate transcriptional activity in combination with the NuRD complex (Popescu et al., 2009). Although exogenously expressed taperin concentrated with CHD7 in HeLa cell nuclei, antibody labeling of endogenous taperin did not accumulate with CHD7 in hair cell nuclei (data not shown). Taperin instead strongly concentrates at the base of stereocilia (Rehman et al., 2010). How might CHD7 and taperin interact despite accumulating in different cellular compartments? Though we did not detect CHD7 in stereocilia by immunofluorescence (data not shown), it is present within the proteome of isolated avian hair cell stereocilia (Shen et al., 2015, Shin et al., 2013), suggesting that CHD7 can shuttle between the hair cell nucleus and stereocilia compartment. Taperin may similarly shuttle, but only be retained within the nucleus in response to cellular stress (Ferrar et al., 2012). Taperin is ideally situated to sense mechanical stress in hair cells, being concentrated at the point where stereocilia pivot during sound stimulation (Flock et al., 1977) and are potentially damaged by noise (Kitajiri et al., 2010). We speculate that taperin and CHD7 may be elements of a mechanism to regulate hair cell gene transcription in response to mechanical stresses experienced by stereocilia during auditory mechanotransduction.

## DISCUSSION

Nanoscale pulldown (NanoSPD) is a powerful new approach for studying multiple aspects of PPIs within live cells, including the assembly of large macromolecular complexes and functional analyses through domain mapping and mutagenesis. Our technique has several important features that complement traditional IP/AP. First, in a nanoscale pulldown PPIs are assayed within the cytoplasm under native conditions, rather than in artificial buffers at non-physiological temperatures. Second, nanoscale pulldowns do not require protein purification, antibodies, or other protein binding reagents to capture macromolecular complexes. This is a key advantage, as it allows the use of folded, full-length proteins that may ordinarily be refractory to purification or difficult to solubilize from a cell. Third, macromolecular complexes are rapidly trafficked and concentrated at filopodia tips, providing biological signal amplification and potentially allowing for low affinity, transient interactions to be detected. This is in contrast to IP/AP, which favors the detection of stable PPIs that remain associated with the affinity matrix during capture and wash steps. Fourth, nanoscale pulldowns have a direct fluorescence readout that allows PPIs to be optically interrogated *in situ* without the additional use of secondary detection techniques (e.g. SDS-PAGE/western blot analysis), which can lack sensitivity if the interaction complex is of low abundance. Finally, in a nanoscale pulldown each cell projects multiple filopodia such that the assay is inherently parallelized to provide multiple independent determinations of an interaction.

We directly visualized bait-prey complexes moving anterogradely along filopodia using TIRFM, demonstrating the core assay principle and the ability of chimeric MY010 molecules to traffic normally (Figure 2). Similar to wild-type MY010, these puncta were in the single molecule regime and challenging to detect (Kerber et al., 2009). Our analysis technique instead focused on the filopodia tips, where the continued flux of MY010 creates a ‘traffic-jam’ of bait-prey complexes. This active accumulation provides fluorescence amplification and allows molecular trafficking to be detected by standard widefield or confocal microscopy. Evaluating fluorescence at the filopodia tips also significantly reduces contaminating background from within the bulk cytoplasm, further contributing to the biologically relevant signal amplification. Combined, these features distinguish nanoscale pulldowns from other live cell fluorescence colocalization techniques that capture freely diffusing prey molecules and tethers them to the plasma membrane, or within the nucleus (Gallego et al., 2013, Herce et al., 2013, Zolghadr et al., 2008).

We describe an analysis pipeline and software tool that allows nanoscale pulldowns to be quantified in a rigorous and unbiased manner. Both analysis techniques use line scans to estimate correlation between bait and prey fluorescence, either along the filopodia shaft (spatial correlation) or at the tips (intensity correlation), to infer the presence of an interaction. Spatial correlation analysis measures relative fluorescence changes, allowing cells to be imaged with different gain and laser parameters. Intensity correlation analysis requires these parameters be kept constant, however it can reveal subtle changes in pulldown efficiency that may relate to binding affinities. The spatial correlation technique had a higher false discovery rate in the short, bulbous filopodia of Sf9 cells, indicating that the analysis technique needs to be matched to the cell type being assayed. Our current analysis pipeline uses manual line tracing of filopodia, which typically requires < 5 minutes of user time per cell analyzed. Recently published algorithms to segment and skeletonize filopodia will help streamline and automate this process (Tsygankov et al., 2014).

Nanoscale pulldowns were able to detect a range of PPIs involving molecular motors (Figure 3), phosphatases (Figure 5), nuclear (Figure 6), membrane-associated (Figure 5), and more generally proteins of unknown function. Despite extensive use in our laboratory, we have yet to encounter a PPI incompatible with nanoscale pulldowns. However, there are two specific examples where the endogenous localization of a prey protein, to either the filopodia tip complex or nucleus, can initially hinder the assay. One approach for assaying nuclear proteins is to mutate/truncate predicted nuclear localization signals (NLS) to prevent the accumulation of prey in the nucleus, where it is segregated from MY010-bait molecules. This was an effective solution for studying the CHD protein superfamily (Figure 6). In general, presenting nuclear and filopodia proteins as bait (i.e. fused to MY010) allows these types of interactions to be successfully tested. If necessary, switching orientation in our current assay requires recloning of the bait and prey plasmids, since we use a chimeric fusion of MY010 and bait molecules. We are now testing MY010 fused to camelid nanobodies engineered to selectively bind either GFP or RFP variants with high affinity (Fridy et al., 2014, Rothbauer et al., 2008). In NanoSPD v2.0, expression of the MY010^GFP-BINDER^ nanobody fusion would capture and selectively traffic all GFP-tagged molecules as the assay bait, and then co-traffic interacting RFP-tagged proteins as prey. Switching assay orientation would then be simply achieved by expressing the MY010^RFP-BINDER^ nanobody fusion, rather than the MY010^GFP^ nanobody variant. This removes the need for generating covalent MY010^BAIT^ fusions, and will allow publicly available repositories of fluorescently tagged expression constructs to be leveraged (Kamens, 2014).

Nanoscale pulldowns measure PPIs in the native cytosolic environment of live cells. Whilst desirable in most situations, this does preclude the intentional variation of buffer conditions and protein concentrations that can be used to estimate equilibrium constants and PPI binding characteristics (Hakhverdyan et al., 2015, Pollard, 2010). Furthermore, like all intracellular based assays, nanoscale pulldowns cannot infer direct binding between bait and prey, since additional native proteins can be recruited from the cytosol. If filopodia could be selectively isolated, then nanoscale pulldowns would be a viable platform to discover these intermediate adapters by mass-spectroscopy or other proteomic methods. Another possible approach would be fusing a promiscuous biotin ligase (BiolD) to the MY010^BAIT^ and using a brief ‘pulse’ of biotin prior to cell lysis and affinity purification to analyze complexes concentrated at the filopodia tips (Roux et al., 2012).

The concept of using intracellular molecular motors to concentrate bait-prey complexes is broadly extensible. We showed that nanoscale pulldowns can be used to study the formation and binding hierarchy within a tripartite protein complex. The introduction of additional wavelength multiplexed fluorescent proteins, or the use of sequential antibody labeling could allow for large numbers of prey to be studied within a complex (Jungmann et al., 2014, Kremers et al., 2011). Additional dark (unlabeled) proteins can also be introduced to examine their effects (positive or inhibitory) upon complex formation (Figure 4). Furthermore, since nanoscale pulldowns are based upon quantitative fluorescence, they can in principle be used with any labeled molecule. An exciting future possibility is labeling nucleic acids in live cells to study the formation of protein-RNA hybrids (Bertrand et al., 1998, Nelles et al., 2016, Paige et al., 2011).

Nanoscale pulldowns are also scalable for high throughput (HT) studies of macromolecular interactions. Whilst HT proteome-wide IP studies have been enormously successful (Hein et al., 2015, Huttlin et al., 2015), the fundamental assay technology still exhibits a significant false discovery rate due to loss of low affinity interactions during capture and wash steps (Gingras et al., 2007). A HT NanoSPD assay could take several forms. For discovery of new interactions, cells would be transfected with pairwise combinations of bait and prey constructs and screened for positive correlation within filopodia. An additional assay format would be to screen small-molecule compound libraries for novel biological targets. Cells would be transfected with a known bait-prey complex and then incubated with a library of candidate small molecules. In both cases, the assay is read optically on a high content screening system with minimal fluid handling required. Combined, these modifications could allow nanoscale pulldowns to accelerate the discovery of novel PPIs and the identification of small molecules that bind to clinically important drug targets.

## ACKNOWLEDGEMENTS

We thank Giovanni Mann and Yasuharu Takagi for insightful discussions, Dennis Drayna, James Sellers and Matthias Machner for critical reading, and Joseph Duda and Alexandra Boukhvalova for technical assistance. We are grateful to John Hammer (NHLBI) for the gift of *Myo5a* and *Mlph* cDNA, and to Stuart Yuspa (NCI) for providing *Clic4* mutant mice, and to the animal care staff at both the NIDCD and the University of Michigan. This work is supported (in part) by the Intramural Research Program of the NIH, NIDCD Z01 DC000039 (to TBF), NIDCD ZIC DC000080 (to TF), and extramural funds NIDCD R01 DC009410 (to DMM).

## AUTHOR CONTRIBUTIONS

**Conception and design:** JEB MB MCD ETB IAB TBF

**Acquisition of data:** JEB MB MCD SMC ELM DCS SMG IAB EAW TF AUR DMM

**Designed analysis and software tool:** JEB

**Analysis and interpretation of data:** JEB MCD MB SMC ELM SMG DCS EAW TF IAB DMM TBF

**Drafted the manuscript:** JEB MB MCD IAB ETB TBF

**Commented and revised the final manuscript:** All authors

### EXPERIMENTAL PROCEDURES

#### Expression Constructs

PCR amplification and cloning were performed using standard methods. All expression constructs were verified by Sanger sequencing. Site-directed mutagenesis (SDM) was performed using QuikChange II SDM Kit (Agilent). Plasmid DNA for transfection was prepared using endotoxin-free purification kits (NucleoBond EF, Clontech). The MY010 heavy meromyosin (HMM) construct generated for the nanoscale pulldown encompasses the ATPase motor, light chain binding sites and a coiled-coil region (residues 1-941; NP_062345) and is not a forced dimer (i.e., MY010^HMM^ was not fused to a leucine zipper or a similar dimerization motif). All expression constructs used in this study are listed in Supplemental Table S1.

#### Animals

All experimental animal procedures were performed in accordance with NIH guidelines, and approved by the respective Institutional Animal Care and Use Committees (IACUC) at the NIDCD (#1263) and the University of Michigan (PR000006244). C57BL6/J mice were obtained from the Jackson Laboratories (Bar Harbor, ME, USA). *Clic4* null mice were described previously (Padmakumar et al., 2012). Conditional null *Chd7* mice were generated by intercrossing mice carrying the conditional floxed *CM7^LoxP^* allele (Hurd et al., 2010), a gene-trapped null *Chd7^GT^* allele (Hurd et al., 2007) and the *FoxG1^Cre^* allele (Hebert et al., 2000) to yield *Chd7^GT/LoxP^:FoxG1+^/Cre^* pups. The *ROSA26^LoxP-STOP-LoxP-ZsGreen1A^* allele was also included on this mutant background (Madisen et al., 2010).

#### Yeast 2-Hybrid Screening

Screening of mouse inner ear and kidney prey libraries (Boeda et al., 2002) was performed by Hybrigenics Services SAS, using a N-terminal LexA fusion with full-length mouse taperin (TPRN, aa 1 – 749, NP_780495) as bait.

#### Measurement of Hearing Function in Mouse

Auditory brainstem responses (ABRs) were measured using a Tucker-Davis Technologies (TDT) hardware (RZ6 Processor) and software (BioSigRZ, v.5.1). Mice were anesthetized by intraperitoneal injection of ketamine (56 mg/kg) and dexdomitor (0.375 mg/kg) and placed on a heating pad connected to a temperature controller (World Precision Instruments) inside a sound-insulated booth (Acoustic Systems). Tone burst stimuli were presented to the test ear at 8, 16, 32, and 40 kHz via a closed-field TDT MF-1 speaker. For each frequency, testing began at 80 dB SPL and decreased in 10 dB steps until the ABR waveform was no longer discernable; testing then proceeded in 5 dB steps. If no response was obtained at 80 dB SPL, testing was performed at the maximum level of 90 dB SPL. ABR thresholds were determined as the lowest stimulus level that elicited repeatable waves.

#### Cell Culture and Transfection

COS-7 (ATCC CRL-1651) and HeLa (ATCC CCL-2) cells were cultured in DMEM supplemented with 10% (v/v) heat-inactivated fetal bovine serum (FBS) (Atlanta Biologicals), GlutaMAX (Thermo Fisher Scientific) and incubated at 37°C, 10% CO_2_. Hela cell transfections were performed with Lipofectamine 3000 (Thermo Fisher Scientific), and COS-7 transfections were performed using the Neon transfection system (Thermo Fisher Scientific) following the manufacturer’s protocol. Transfected cells were incubated for 12-24 hours before seeding on fibronectin-coated (10 μg/ml, Sigma) glass-bottom culture dishes (#1.5, MatTek Corporation). Cells were incubated for another 6-24 hours prior to fixation with 4% paraformaldehyde (PFA) for 15 mins. For live-cell imaging, transfected cells were incubated for 1-3 hours after seeding and then exchanged into FluoroBrite DMEM (Thermo Fisher Scientific) supplemented with GlutaMAX.

Sf9 (*Spodoptera frugiperda*) cells (Thermo Fisher Scientific) were cultured in HyClone SFX media (GE Healthcare) supplemented with GlutaMAX and heat-inactivated 2.5% (v/v) FBS (Atlanta Biologicals) and maintained at 27°C in a shaking incubator. For transfection, DNA was diluted into phosphate buffered saline (PBS) at a 1:12 ratio (w/w) with polyethylenimine (1 mg/ml, PEI Max, #24765 Polysciences Inc.), incubated for 15 mins at RT, before adding the complexed DNA to cells in suspension culture. 48 hours post-transfection, cells were seeded onto glass-bottom dishes (#1.5, MatTek Corporation) for 1-3 hours at 27°C prior to fixation with 4% PFA for 15 mins.

Mammalian and insect cells were co-labeled with either Atto 390 conjugated phalloidin (Sigma) or Alexa Fluor conjugated phalloidin to visualize the filamentous actin cytoskeleton, and 4′,6-diamidino-2-phenylindole (DAPI) / Hoechst 33342 (Thermo Fisher Scientific) to stain the nucleus.

#### Organotypic Cochlear Culture and Biolistic Transfection

Biolistic transfection of plasmid DNA into cochlear explants was performed as described previously (Belyantseva, 2009). Briefly, organ of Corti explants were dissected from cochleae of C57BL/6J mice at P3, and cultured in DMEM supplemented with 7% FBS (Thermo Fisher Scientific) on glass-bottomed culture dishes (MatTek) coated with rat tail collagen (Thermo Fisher Scientific) at 37°C, 10% CO_2_. After 24 hours *in vitro*, cochlear explants were transfected with plasmid DNA encoding EGFP-CLIC4 using a Helios Gene Gun (Bio-Rad). Gold microcarriers (1.0 μm, Bio-Rad) were coated with plasmid DNA according to the manufacturer’s standard protocol. Cochlear explants were cultured for a further 24 hours post-transfection and then fixed for 2 hours at room temperature in 4% PFA and processed for immunofluorescence as described below.

#### Cochlear Immunocytochemistry

Mice were euthanized and temporal bones fixed for 2 hours in 4% PFA. Micro-dissected samples were permeabilized for 30 mins in 0.5% Triton X-100, and blocked for 1 hour with 2% bovine serum albumin (Roche) and 5% normal goat serum (Sigma) diluted in PBS. Samples were incubated for 2 hours to overnight in blocking solution containing primary antibodies recognizing TPRN (#HPA020899, Sigma) or CHD7 (#6505, Cell Signaling Technology). After washing in PBS, primary antibodies were detected using Alexa Fluor-conjugated secondary antibodies (Thermo Fisher Scientific). Samples were co-labeled with Alexa Fluor 633 or rhodamine-conjugated phalloidin to visualize the actin cytoskeleton, in addition to the nuclear stain Hoechst 33342 prior to mounting in Prolong Gold Antifade Reagent (Thermo Fisher Scientific).

#### Fluorescence Microscopy

For live cell imaging, an inverted microscope (Axiovert 200M, Zeiss) was used to perform objective-based total internal reflection microscopy with a lase*r*-based illuminator (TIRF3, Zeiss) and a 100x 1.46 N.A. oil objective (alpha-Plan Apochromat, Zeiss). EGFP and mCherry fluorescence was sequentially excited through a laser clean-up filter (ZET488/561x, Chroma) and a 2 mm dichroic mirror (ZT488/561rpc, Chroma) with 488 and 561 nm laser lines, respectively. Epifluorescence was collected through a laser-blocking filter (ZET488/561m-TRF, Chroma) and a dual-camera adaptor (Zeiss). Split channels were emission filtered for EGFP (ET525/50m, Chroma) and mCherry (ET610/75m), respectively, and captured on individual EM-CCD cameras (Evolve 512, Photometrics) controlled by Zen software (Zeiss). Dual camera EGFP and mCherry data were re-registered with MATLAB (Mathworks) using fluorescent beads as fiducial markers (Tetraspeck, Thermo Fisher Scientific). Cells were maintained at 37°C in a humidified 10% CO2 environment during all live-cell image acquisitions. Images were captured at 1-2 frames per second.

Fixed cells were imaged using a 63x 1.4 N.A. oil objective (Plan-Apochromat, Zeiss) and an Axiovert 200M inverted microscope with a confocal scan head (LSM780, Zeiss). Bait and prey signals were captured sequentially to eliminate spectral bleed-through. The LSM780 GaAsP spectral detector was set to capture 500 – 530 nm for EGFP and 570 – 620 nm for mCherry/tdTomato.

#### Analysis of Fluorescence Co-Localization in Filopodia

Image analysis was performed on confocal images of fixed cells to quantify bait and prey fluorescence at filopodia tips. Line scans along individual filopodia were prepared in ImageJ (http://rsbweb.nih.gov/ij/) using the bait and/or phalloidin (F-actin) fluorescence signal as a guide. Prey fluorescence was kept hidden during line tracing to avoid any selection bias. Filopodia suitable for analysis projected > 2 μm away from the cell body, to allow for comparisons between the filopodia tip and shaft. A maximum of 10 filopodia were sampled per cell, to avoid any single cell from dominating the overall dataset. For each interaction we typically examined 150 filopodia pooled from three independent experiments. Bait and prey fluorescence line scan values were exported from ImageJ into a tab-delimited text file, and all further data analysis was performed in the MATLAB (Mathworks). Compiled binary distributions, source code and a detailed guide to using this software tool can be found in the Supplemental Materials.

Within MATLAB, the global maximum of the MY010^BAIT^ signal was used to determine the position of the filopodia tip. This approximation was valid given the robust accumulation of MY010 at filopodia tips, even when fused to different bait domains. Line scan data were then screened for basic quality criterion, including the presence of a robust accumulation of the MY010^BAIT^ molecule. Filopodia were excluded if the peak bait fluorescence at the tip did not exceed the average bait fluorescence along the shaft by more than three standard deviations. Filopodia were also excluded if the full-width half maximum (FWHM) of the tip bait fluorescence was too large, as this indicated the bait was not sufficiently concentrated at the tip. Filopodia line scans passing these quality criteria were analyzed further by one of two algorithms.

#### Spatial Correlation Analysis

In the presence of an interaction, we assume that the distribution of prey fluorescence along an individual filopodium will mirror the bait fluorescence. Pearson’s coefficient (*r*) was used to quantify the correlation between bait and prey fluorescence data along individual filopodia. The *r* coefficient yields values −1 ≤ *r* ≤ +1, where *r* = +1 indicates perfect linear correlation, *r* = −1 indicates perfect linear anti-correlation and *r* = 0 indicates no correlation. Bait and prey fluorescence values should be perfectly correlated (i.e. *r* = +1) in the presence of an interaction, and uncorrelated (i.e. *r* = 0) in the absence. The use of a global threshold to determine the significance of any given bait-prey correlation is insensitive, and will likely discard weak correlations that are potentially of biological significance. Instead, we estimated the likelihood that the observed correlation could have occurred by random chance. Computationally, this was implemented by keeping the bait line scan data constant, whilst continually randomizing the data order of the prey line scan and measuring the resulting *r*-values (Costes et al., 2004, Lifshitz, 1998). Bootstrapping over a large number of trials assembles an exact probability distribution of all possible correlations. A refinement to this methodology is to perform the randomization in blocks that approximate the point spread function of the microscope. This preserves inherent correlation that is present in adjacent pixels (Costes et al., 2004). This exact probability distribution was then used to calculate the likelihood (P value) that any observed correlation occurred by chance. A filopodium was scored as a positive bait-prey interaction if: 1) the calculated P value < 0.01, and 2) the prey intensity at the filopodia tip exceeded the filopodia shaft signal by three standard deviations. For each experimental determination, an interaction index is calculated as the ratio of correlated filopodia to the total number of filopodia assayed.

#### Intensity Correlation Analysis

In this alternate algorithm, which is strongly recommended for Sf9 cells, we test for the presence of a global correlation between bait and prey fluorescence intensities at filopodia tips. These two quantities are expected to be positively correlated in the presence of an interaction, with the prey fluorescence being dependent upon the bait fluorescence. In this approach, statistical significance is determined globally, and not on a per-filopodia basis (see spatial analysis above). Line scan data from ImageJ were imported into MATLAB and filtered as described above. The maximum bait and prey fluorescence measured at the filopodia tip is then exported for each filopodia. Paired bait and prey fluorescence values were plotted on an X-Y scatter chart, with each data point representing one filopodium. Since this algorithm depends upon absolute fluorescence values, it is critical that imaging conditions (objective, gains/exposures, illumination intensity) are kept constant between independent determinations.

#### Statistical Analyses

All statistical testing was performed in Prism (v7.0, GraphPad). Datasets were analyzed for normality using the Shapiro-Wilk test. Comparisons between two treatment groups were made using Student’s *t*-test (assuming unequal standard deviation) for normally distributed data, and by non-parametric Mann Whitney *U* test otherwise. For multiple comparisons, one-way analysis of variance (ANOVA) was used with Tukey’s post-hoc test. All data are mean ± standard deviation, and from three independent experimental determinations, unless otherwise stated. Statistical significance is indicated as: (*) P < 0.05, (**) P < 0.01, (***) P < 0.001, (****) P < 0.0001.

---

**Figure S1.**
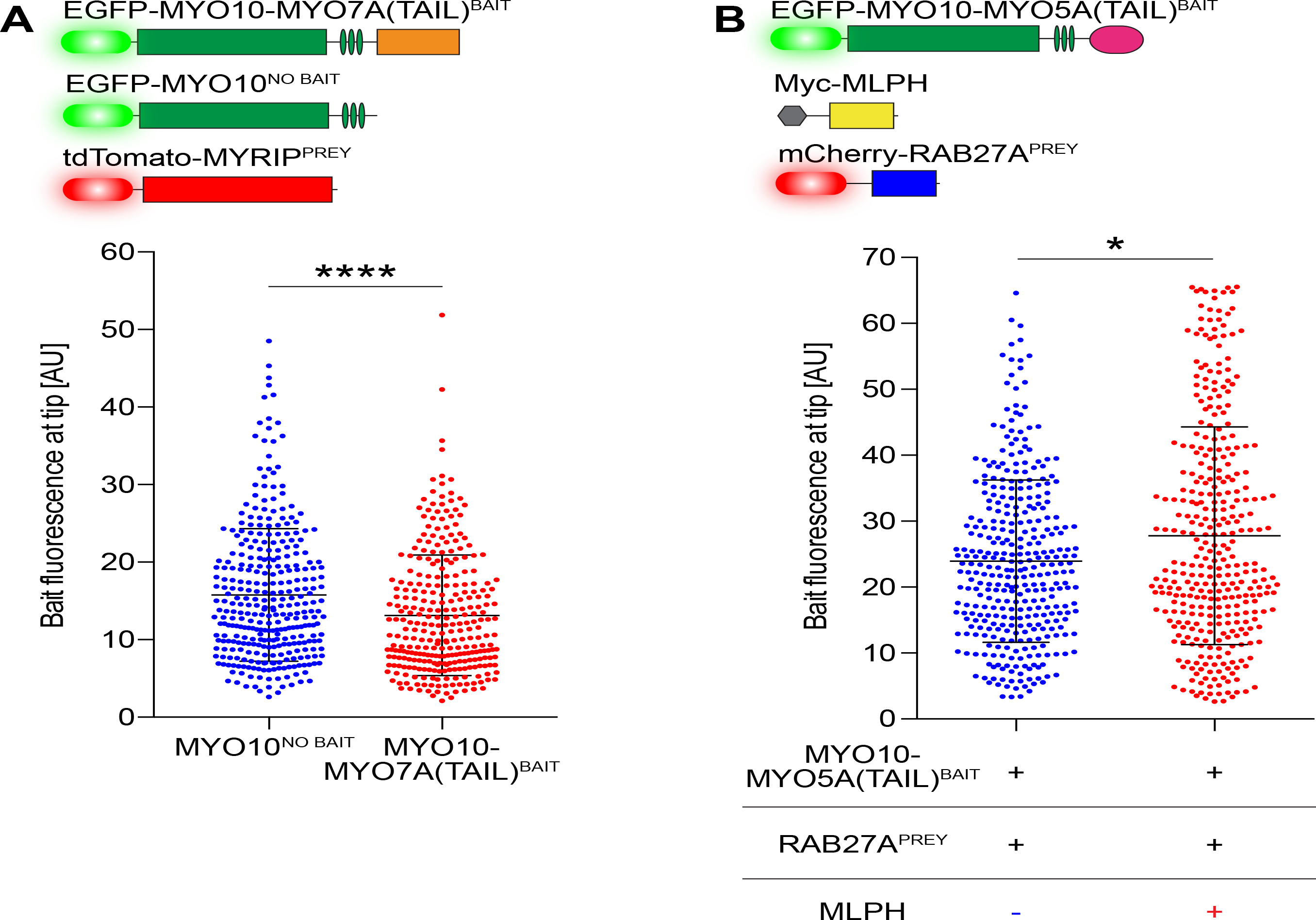
**Fusion of bait molecules to MYO10 does not interfere with filopodia trafficking (A)** Comparison of bait fluorescence intensities at filopodia tips of Sf9 cells expressing MY010^NO BAIT^ + MYRIP^prey^, or MY010-MY07A(TAIL)^bait^ + MYRIP^prey^. Though bait fluorescence is statistically different (Mann-Whitney *U* test), the effect size is minimal indicating that the MY010 motor domain traffics equally, with (red) or without (blue) a bait molecule attached. **(B)** Bait intensity of MY010-MY05A^BAIT^ at Sf9 filopodia tips is unaffected by the absence (blue) or presence (red) of myc-MLPH, indicating no systematic change in bait molecule trafficking (Mann-Whitney *U* test). Supplementary Figure S1A accompanies Figure 3, Supplementary Figure S1B accompanies Figure 4.

**Figure S2.**
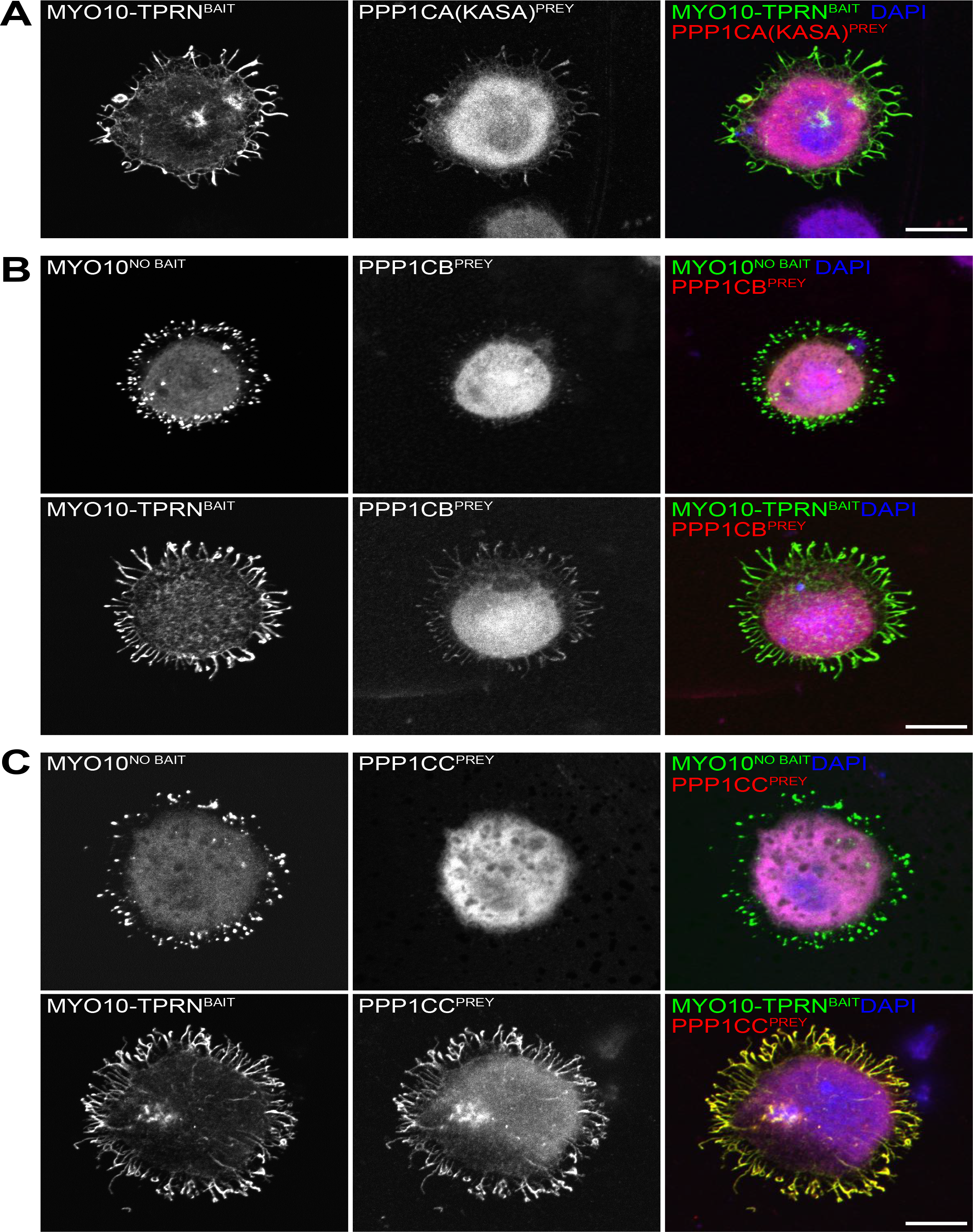
**Taperin selectively interacts with the catalytic subunits of PP1 (A)** Two point mutations in a conserved motif (KISF → KASA) within taperin disrupt its interaction with PPP1CA. **(B)** PPP1CB does not significantly interact with taperin. **(C)** Taperin interacts with PPP1CC. All panels: Top row, control (bait omitted); bottom row, bait + prey. Sf9 cell nuclei are labeled with DAPI. Scale bars, 10 μm. Supplementary Figure S2 accompanies Figure 5.

**Figure S3.**
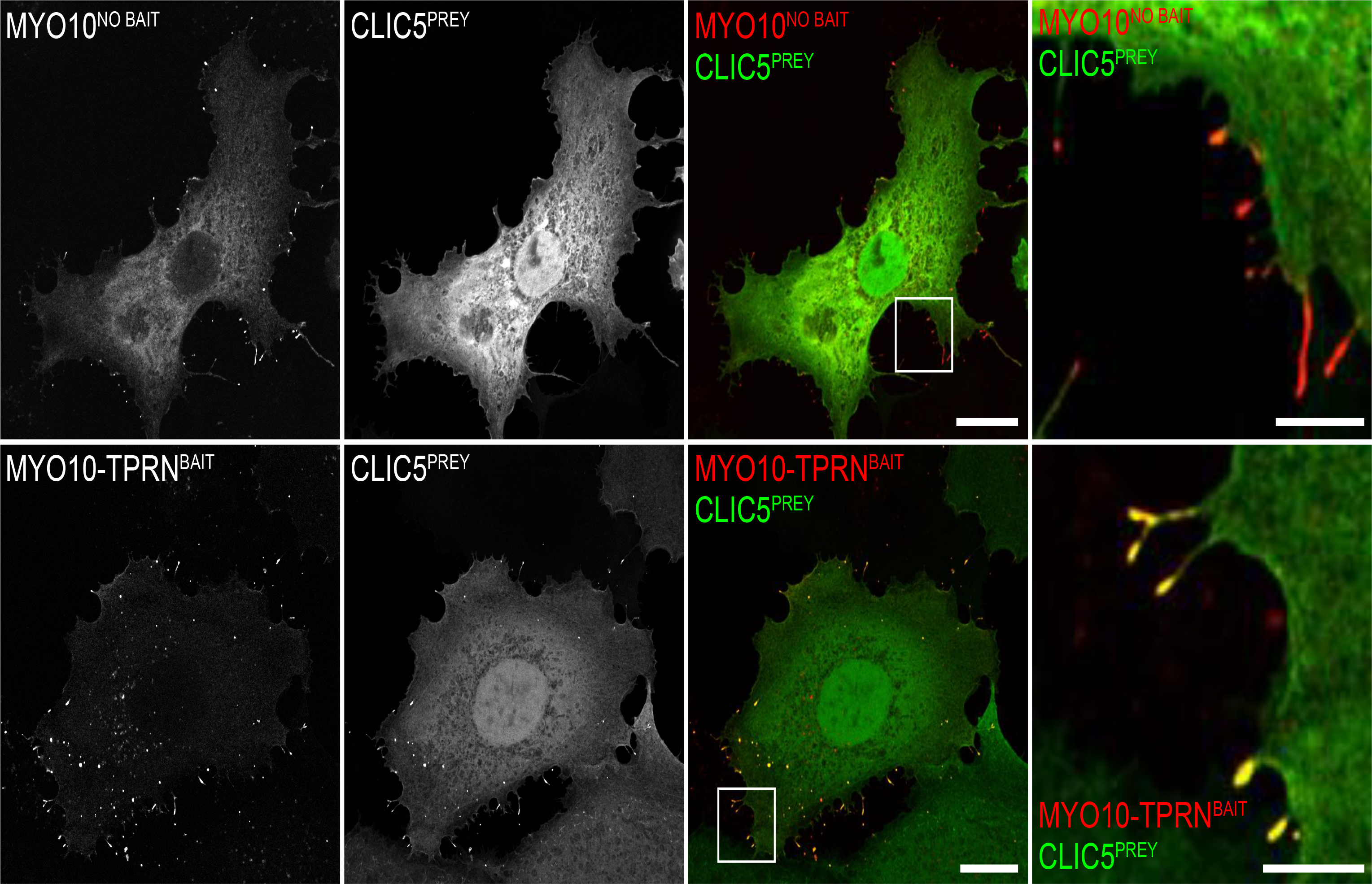
**Taperin interacts with CLIC5** CLIC5 binds taperin. CLIC5^PREY^ was co-expressed with either MY010^NO BAIT^ (control, top row) or MY010-TPRN^BAIT^ (query, bottom row). Boxed insets are shown magnified (right column). F-actin is labeled with rhodamine phalloidin (red). Scale bars, 20 μm; 5 μm in magnified insets. Supplementary Figure S3 accompanies Figure 5.

**Figure S4.**
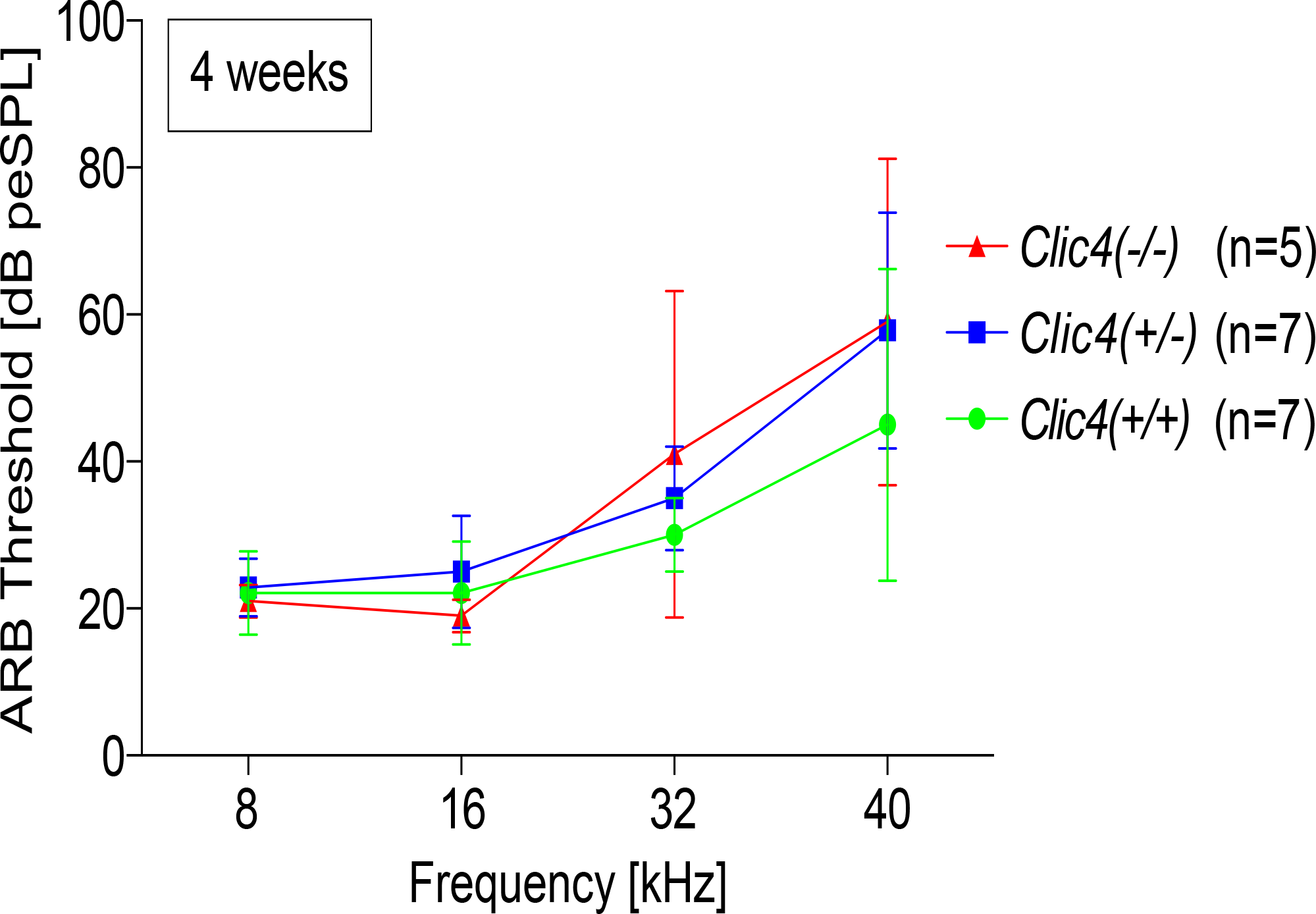
***Clic4* mutant mice have normal hearing** Auditory brainstem response (ABR) thresholds of 4 week old mice measured at 8, 16, 32 and 40 kHz. No significant difference in ABR thresholds was detected between wild-type *Clic^4+/+^* (green), heterozygous *Clic4^+/−^* (blue) and homozygous *Clic4^−/−^* mutant (red) mice. n = 5 – 7 mice per group. Supplementary Figure S4 accompanies Figure 5.

**Table S1.**
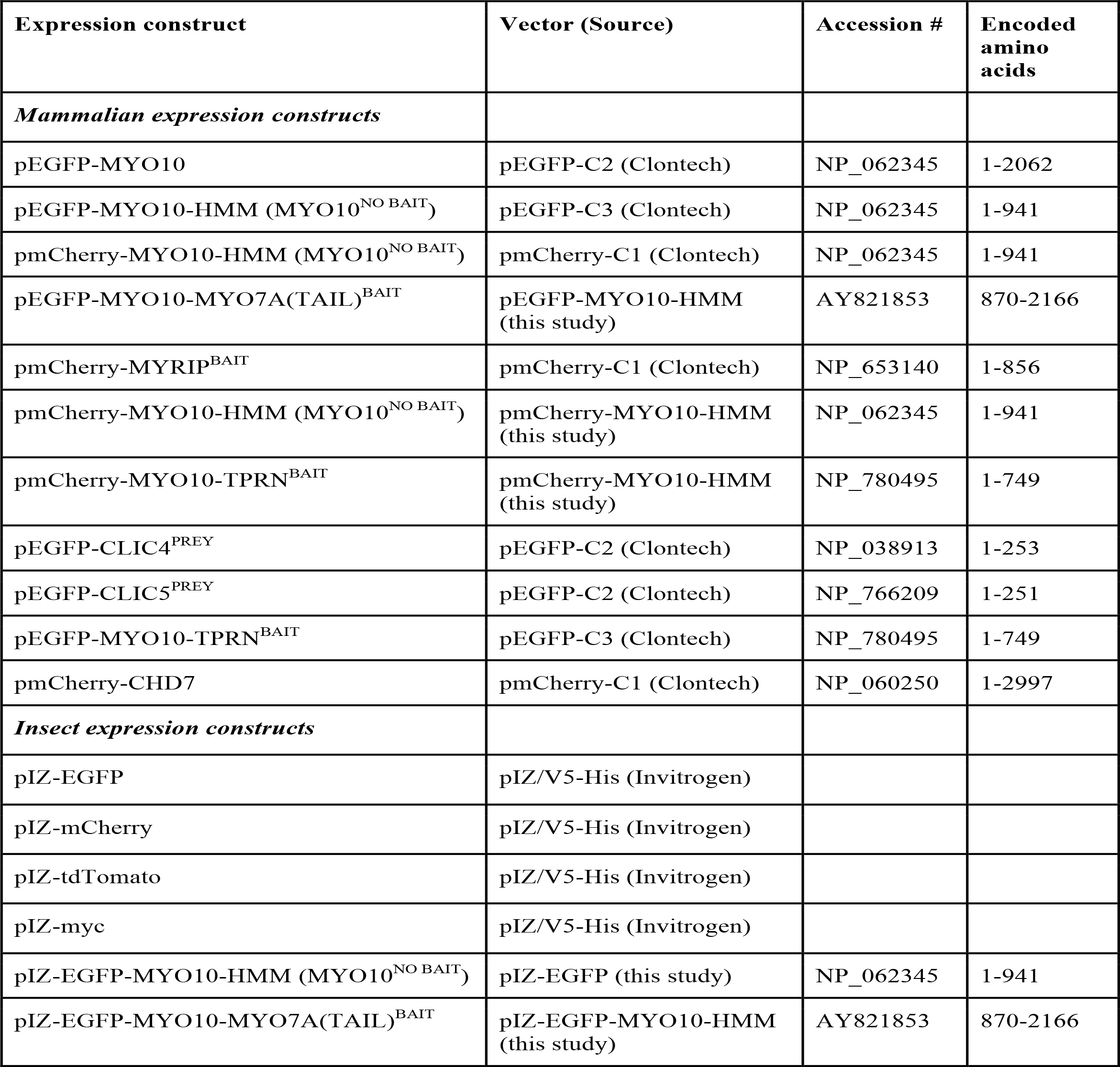
Plasmids used in this study

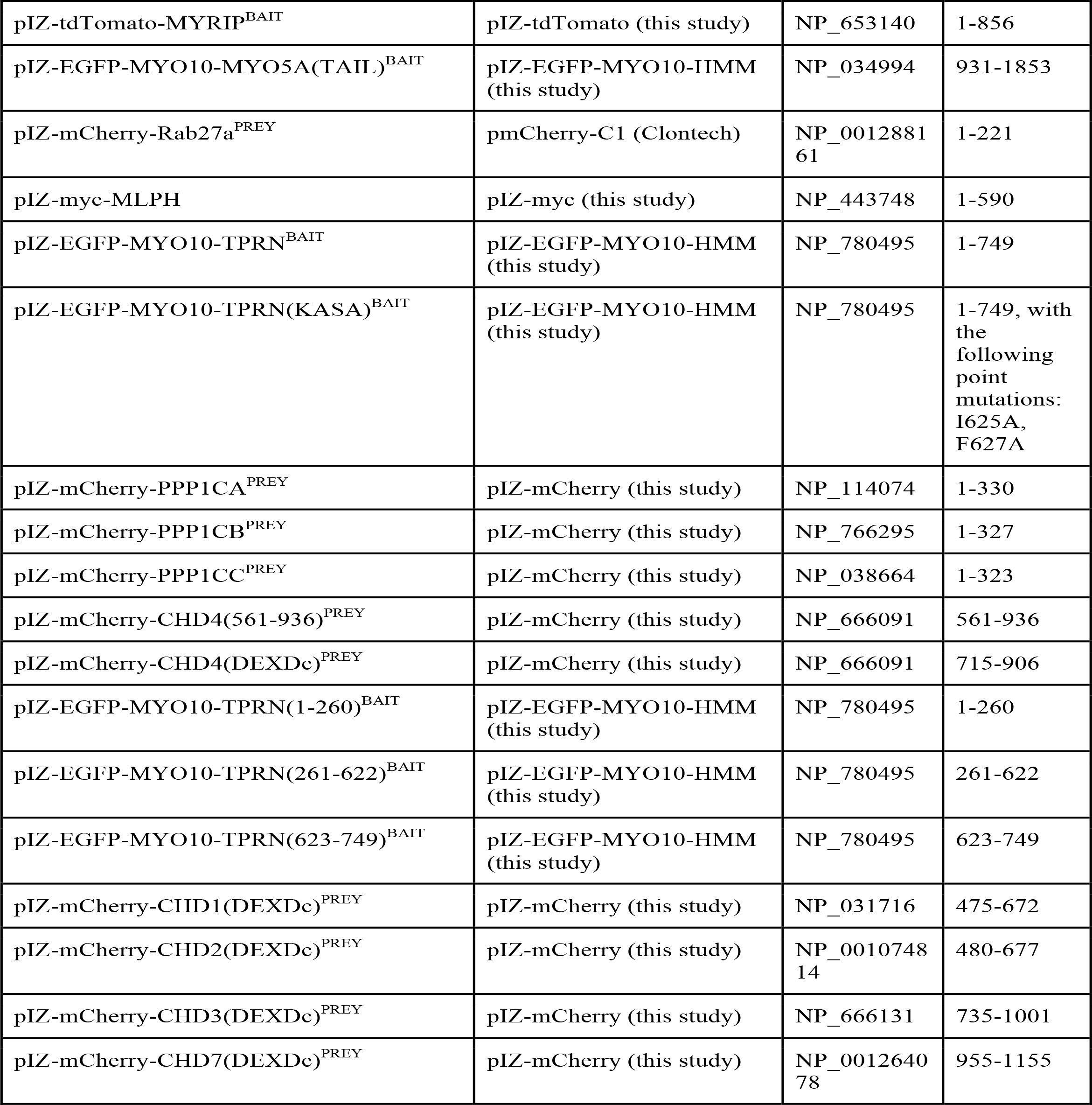

**Table S2.** Yeast 2-hybrid interactions identified with taperin Results from mouse inner ear Y2H screen

| Gene name | # of clones | Confirmed by nanoscale pulldown? |
| --- | --- | --- |
| Chromodomain helicase DNA binding protein 4 (*Chd4*) | 16 | **Yes** |
| Protein phosphatase 1 catalytic subunit alpha (*Ppp1ca*) | 5 | **Yes** |
| Chromodomain helicase DNA binding protein 3 (*Chd3*) | 3 | **Yes** |
| Chloride intracellular channel 4 (*Clic4*) | 1 | **Yes** |
| Hes related family bHLH transcription factor with YRPW motif-like (*Heyl*) | 3 | Not tested |
| Chloride intracellular channel 1 (*Clic1*) | 2 | Not tested |
| Fatty acid synthase (*Fasn*) | 2 | Not tested |
| Protein disulfide isomerase family A member 4 (*Pdia4*) | 2 | Not tested |
| Germ cell-less spermatogenesis associated 1 (*Gmcl1*) | 1 | Not tested |
| Lysine acetyltransferase 5 (*Kat5*) | 1 | Not tested |
| Actin binding LIM protein 1 (*Ablim1*) | 1 | Not tested |
| Zinc finger and BTB domain containing 44 (*Zbtb44*) | 1 | Not tested |

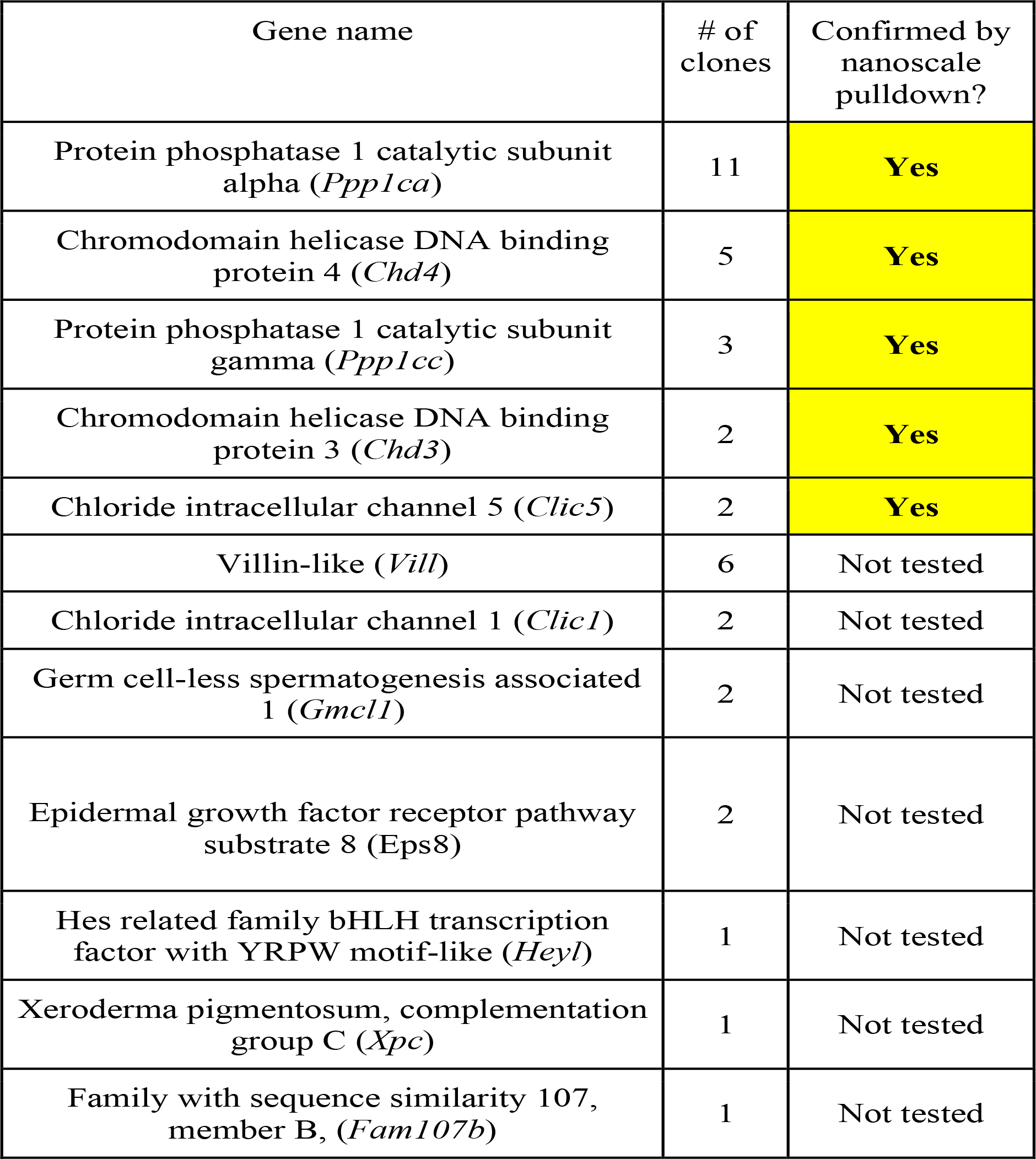

